# Membrane binding and pore formation is Ca^2+^-dependent for the *Clostridioides difficile* binary toxin

**DOI:** 10.1101/2023.08.18.553786

**Authors:** Dinendra L. Abeyawardhane, Spiridon E. Sevdalis, Kaylin A. Adipietro, Raquel Godoy-Ruiz, Kristen M. Varney, Izza F. Nawaz, Alejandro X. Spittel, Richard R. Rustandi, Vitalii I. Silin, Amedee des Georges, Mary E. Cook, Edwin Pozharski, David J. Weber

## Abstract

The *C. difficile* binary toxin (CDT) enters host cells via endosomal delivery like many other ‘AB’-type binary toxins. In this study, the cell-binding component of CDT, termed CDTb, was found to bind and form pores in lipid bilayers upon depleting free Ca^2+^ ion concentrations, and not by lowering pH, as found for other binary toxins (i.e., anthrax). Cryoelectron microscopy, nuclear magnetic resonance spectroscopy, surface plasmon resonance, electrochemical impedance spectroscopy, CDT toxicity studies, and site directed mutagenesis show that dissociation of Ca^2+^ from a single site in receptor binding domain 1 (RBD1) of CDTb is consistent with a molecular mechanism in which Ca^2+^ dissociation from RBD1 induces a “trigger” via conformational exchange that enables CDTb to bind and form pores in endosomal membrane bilayers as free Ca^2+^ concentrations decrease during CDT endosomal delivery.

**One-Sentence Summary:** Ca^2+^ dissociation and unfolding of a receptor binding domain in CDTb activates pore-formation by the *C. difficile* binary toxin.

## Main Text

The Centers for Disease Control and Prevention identified *Clostridioides difficile* infection (CDI) as one of the five most urgent bacterial threats to address in the United States. *Clostridioides difficile* is a gram-positive anaerobic bacterial pathogen responsible for a high-risk nosocomial disease prevalent among patients undergoing prolonged antibiotic treatments and/or cancer therapy (*1*). Reduced levels of symbiotic gut microbiota allows for dominance of *C. difficile* bacteria leading to severe diarrhea and pseudomembranous colitis (*2*). Treatment of severe CDI is difficult because therapeutic options are available only for the large clostridial toxins, termed TcdA (Toxin A) and TcdB (Toxin B) (*3, 4*), which is a problem since CDIs associated with hypervirulence and high recurrence rates often have a third toxin termed the *C. difficile* transferase (CDT) for which no treatment options are available (*5*). CDI studies completed *in vivo* demonstrate that inhibiting the binary toxin (CDT), in addition to the large clostridial toxins (TcdA, TcdB), must be achieved to protect from morbidity associated with CDT-containing strains of CDI (*6*).

CDT enters host cells via endosomal delivery like many other ‘AB’-type binary toxins including *Clostridium perfringens*, *Clostridium spiroforme*, *Clostridium botulinum*, and *Bacillus anthracis* (*7–9*). Upon cellular entry via CDTb, the 47.4 kDa CDTa ribosyltransferase component of the binary toxin modifies intracellular actin resulting in its depolymerization, cytoskeleton destruction, and rapid host cell death. CDTa is composed of an N-terminal domain that binds to CDTb and a C-terminal catalytic domain (*10*). Pro-CDTb (99 kDa) undergoes proteolytic cleavage by trypsin/chymotrypsin to form active CDTb subunits (75 kDa). Under soluble conditions and at millimolar Ca^2+^ ion concentrations found in the extracellular environment, CDTb subunits can oligomerize into two stable “dumbbell-shaped” dimer of heptamers (*11*). The functional importance of the di-heptamer assemblies is somewhat controversial, but they are hypothesized to stabilize the active 75 kDa subunit of CDTb and “protect” it from an immune response and/or continued extracellular proteolytic cleavage prior to host cell binding (*11, 12*). The di-heptameric structures are distinct and include a symmetric (^Sym^CDTb) and an asymmetric (^Asym^CDTb) form (*11*). ^Asym^CDTb has one heptamer unit that forms an extended β-barrel originating from its heptameric core while ^Sym^CDTb lacks the β-barrel extension and displays two identical heptamer units. The di-heptamer interfaces for both ^Asym^CDTb and ^Sym^CDTb involve unique “donut-like” configurations comprised of receptor binding domain 2 (RBD2). ^Sym^CDTb is in an “open” pore state and ^Asym^CDTb is in a “closed” state (*11, 12*). The open/closed terminology reflects an inner diameter of what is called the χπ-gate, which is composed of side chains from phenylalanine residues (F455) that originate from the heptamerization domain 2 (HD2) of each subunit of CDTb. For hypervirulent CDI, it is hypothesized that the toxic CDTa component of CDT traverses from endosomes into the host cytosol in an unfolded conformation via the χπ-gate of CDTb in a manner analogous to other AB-type binary toxins (*3*). However, the molecular mechanism for CDTb host cell interaction(s) and CDTa delivery is not elucidated fully despite the availability of several high-resolution structures for CDTa, CDTb, and other relevant CDT binary complex (*10, 11, 13, 14*).

Based on mechanisms established for anthrax toxin, it was anticipated that CDTb would undergo a conformational shift facilitated by entry into an acidic pH found in early endosomal compartments that would allow for transport of the catalytic component, CDTa, into the cytosol (*15*). However, lowering pH alone was <u>not</u> sufficient for CDTb to bind membrane and form pores, as found for the cell-binding component of anthrax, termed the protective antigen (PA). As there is a significant drop in free Ca^2+^ ion concentration upon entering endosomes from the extracellular environment (*3, 16*), we next considered whether CDTb binding to membranes and pore formation would respond to lowering free Ca^2+^ ion concentrations. Surprisingly, Ca^2+^-dissociation from a site in the CDTb receptor binding domain 1 (RBD1) was indeed found to promote CDTb binding to lipid bilayers and pore formation. Studies next of the structure and dynamic properties of CDTb demonstrated that removal of Ca^2+^ from the RBD1 site induced conformational exchange throughout the entire RBD1 domain, which “triggered” (i) di-heptamer to heptamer transition, (ii) CDTb binding to lipid bilayers, and (iii) CDTb-dependent pore formation. Whether Ca^2+^-dissociation, pH-change(s), and/or a combination of these physiologically relevant gradients contribute to endosomal delivery of other AB binary toxins is important to consider now, based on the findings reported here for CDTb.

### Membrane binding and pore formation by CDTb, the cell-binding component of CDT

A fundamental question regarding host engagement was considered regarding whether the cell-binding component of CDT could insert into lipid bilayers and form pores, as found for other members of the AB binary toxin family, such as the protective antigen (PA) of anthrax (*17, 18*). The question was addressed using a combination of surface plasmon resonance and electrochemical impedance spectroscopy (SPR/EIS). A custom-built experimental platform was used to measure CDTb adsorption/desorption at the biomimetic tethered bilayer lipid membrane via a change of SPR signal while measurement of electrical resistance of the bilayer via EIS was recorded in parallel (*19, 20*). Previous work showed that cellular uptake of *C. perfringens* iota toxin, *C. botulinum* C2 toxin, and anthrax PA are all facilitated by acidification, so it was anticipated that CDTb may undergo a conformational change facilitated by acidic pH upon entry into early endosomal compartments to facilitate transport of the catalytic component, CDTa, into the cytosol (*15, 21, 22*). Therefore, we lowered the pH to match values typically found in endosomes (∼pH 5.5) and examined whether CDTb insertion into membrane and pore formation could be observed using EIS upon binding to POPC-based tethered bilayer lipid membrane, which is the major constituent of endosomal membrane (*23*). However, unlike what was found previously for other toxins, and verified here for anthrax PA (Fig. S1) (*24, 25*), CDTb did not bind lipid bilayers or form pores in membrane-like vesicles upon lowering pH to acidic values.

Upon endosomal compartmentalization, Ca^2+^ ion concentrations also decrease from values in the millimolar range, as found in extracellular space, to low micromolar levels as measured in endosomes (*16*). We therefore examined whether lowering free Ca^2+^ concentration enabled CDTb binding to lipid bilayers to elicit CDTb-dependent pore formation. This hypothesis was intriguing considering a Ca^2+^-binding site was discovered within the RBD1 domain of CDTb (Fig. S2) and that Ca^2+^ dissociation from RBD1 induced pronounced conformational exchange (*11*). To test this hypothesis, we dialyzed the CDTb protein against a buffer containing 2 mM of EDTA and EGTA to decrease free Ca^2+^ ion concentrations as needed to induce conformational exchange in RBD1. As demonstrated by simultaneous increases in SPR signal and electrical conductance by EIS (Fig. 1B), Ca^2+^ depleted CDTb (CDTb^(-Ca)^) was found to adsorb into the phospholipid bilayers and form pores that generates an influx of ions. The membrane binding data together with the increase in EIS signal signifies evidence for channel formation upon CDTb addition, at depleted Ca^2+^ ion concentrations, and supports the hypothesis that Ca^2+^ ion depletion is required to trigger membrane insertion of the CDTb translocase *in vivo*. Whereas the analogous experiments done in the presence of mM Ca^2+^ ion concentrations showed negligible changes in SPR signal and electrical conductance, indicating that neither membrane binding nor pore-formation occurs at millimolar Ca^2+^ concentrations as found outside the cell (Fig. 1A).

**Fig. 1.**
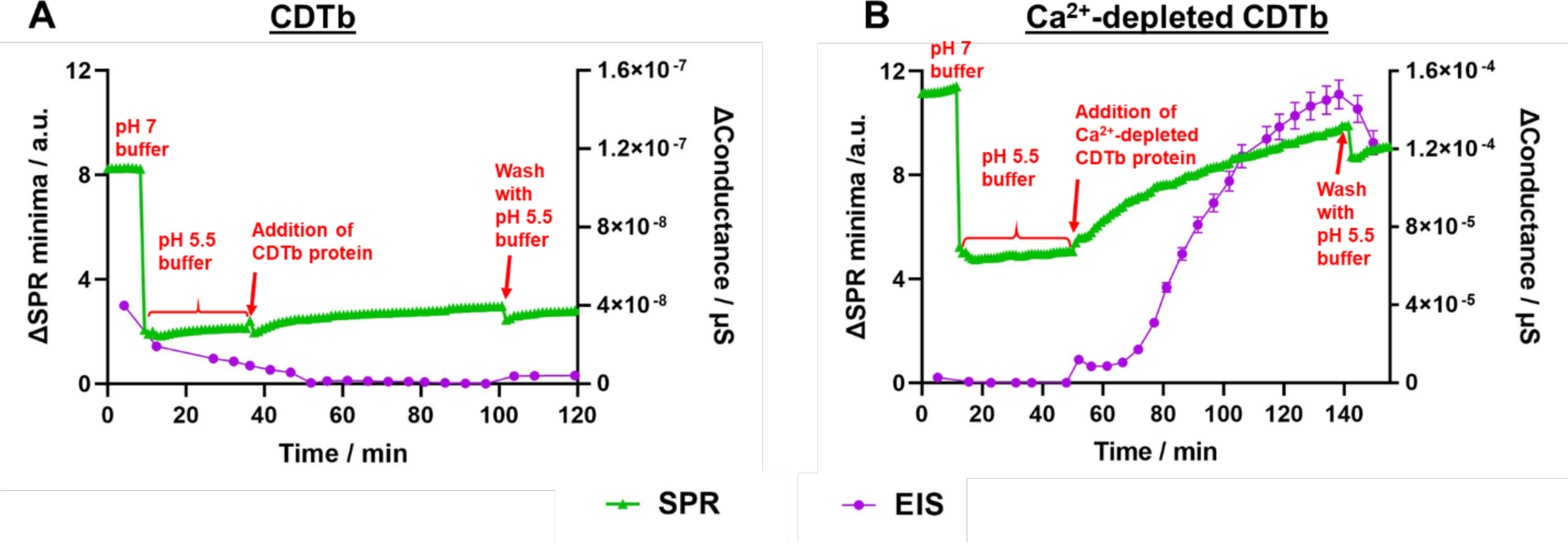
Membrane interactions of CDTb variants at pH 5.5 studied by surface plasmon resonance (SPR, green triangles) coupled to electrochemical impedance spectroscopy (EIS, purple spheres) collected in the (A) absence or (B) presence of the calcium ion chelator EGTA (see Methods).

### CryoEM structures of CDTb in a Ca^2+^-depleted state

Structural studies showed previously that there are at least three Ca^2+^-binding sites on CDTb at mM Ca^2+^ levels as found in the extracellular environment (*11*). This includes two high-affinity Ca^2+^ ion sites in the N-terminal heptamerization domain of CDTb (HD1), which are conserved in anthrax toxin PA and iota toxin Ib, and are essential for binding of the enzymatic component of the binary toxin (*26, 27*). The third Ca^2+^-binding site in CDTb is weaker and involves coordinating residues from within RBD1 of CDTb (Figs. S2, S3). The RBD1 Ca^2+^ binding site was of interest, as it induces conformational exchange throughout the RBD1 domain as Ca^2+^ concentrations are decreased (*11*). Thus, it was hypothesized that Ca^2+^ unbinding from this same RBD1 site, which coincides with an increase in its conformational dynamics, could “trigger” CDTb membrane binding and pore formation.

To test this “Ca^2+^ unbinding” hypothesis further, we solved the structure of wild-type CDTb sample that was dialyzed versus buffer containing 2 mM of EDTA/EGTA using single-particle cryo-electron microscopy (Fig. 2). While these conditions were sufficient to remove Ca^2+^ from the weaker RBD1 site, the higher-affinity dual Ca^2+^ ion binding sites in the HD1 domain of CDTb remained intact (Fig. S4). That Ca^2+^ binding to HD1 was not disrupted by this sample preparation was consistent with reports that disrupting Ca^2+^-binding to HD1 required harsh denaturing conditions for their removal (*28*). Thus, the milder dialysis conditions used here to remove Ca^2+^ from RBD1 did not affect Ca^2+^ binding to HD1 nor did such conditions disrupt the structural integrity of the heptamerization core of CDTb (Fig. 2).

**Fig. 2.**
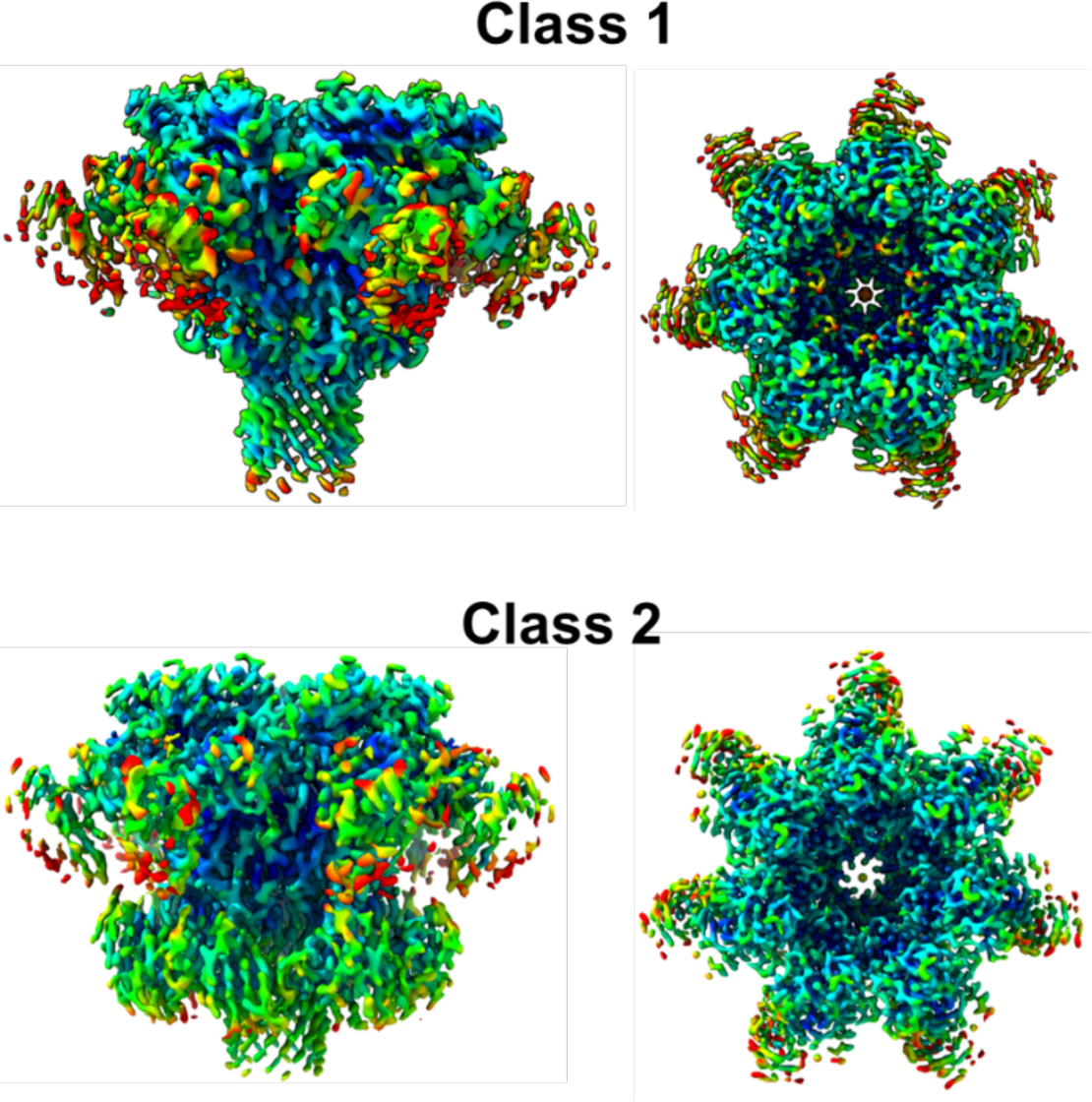
Local resolution in electron density maps of CDTb^(-Ca)^ conformations; RBD2 distorted structure (class 1) and the heptamer with RBD2 intact (class 2). The difference in the degree of RBD2 is distinguishable by side and top projections of each class.

Initial 2D classification for Ca^2+^-depleted CDTb revealed the presence of a single heptamer unit, and subsequent 3D refinement found two distinct structural moieties derived from the “pore-forming” conformation of CDTb (Fig. 2). The difference between the two structures was the degree of flexibility observed for RBD2. An RBD2 distorted structure (class 1) was solved at a resolution of 3.06 Å and the heptamer structure with RBD2 intact (class 2) was solved at 3.28 Å resolution (Fig. 2, Fig. S3). While the elongated β-barrel was not observed in either structure, partial β-barrel formation was detected. As expected, the RBD1 domains in both structures were mobile as is consistent with Ca^2+^ removal from the RBD1 site when this domain was isolated (9). It is interesting that the two Ca^2+^-depleted structures solved (classes 1, 2) both resemble the “prepore states” for CDTb (*14*).

### Structure/function studies of a Ca^2+^-binding mutant of CDTb

We next examined CDTb constructs that eliminated Ca^2+^-binding to RBD1 via a D623A/D734A double mutation (Fig. S2). We first showed by 2D ^1^H-^15^N heteronuclear single quantum coherence (HSQC) NMR that the D623A/D734A double mutation was sufficient to eliminate Ca^2+^ binding to ^15^N-labeled RBD1^D623A/D734A^, as engineered (^Ca^K_D_ >10 mM), at both pH 5.0 and 7.0 (Figs. S5 and S6). In the absence of Ca^2+^, the 2D ^1^H-^15^N HSQC data for RBD1^D623A/D734A^ exhibited significant exchange broadening at both acidic and neutral pH values (pH 5.0 and 7.0) as was anticipated. Whereas upon 10 mM Ca^2+^ addition, no significant changes in the HSQC spectra were observed for RBD1^D623A/D734A^ at either pH value, as compared to wild-type protein, which showed yielded a spectrum that was consistent with a folded RBD1 domain (Fig. 3). These NMR data indicate the RBD1^D623A/D734A^ no longer bound Ca^2+^, even at mM concentrations. Whereas, Ca^2+^-binding and folding of this domain was observed for ^15^N-labeled RBD1^WT^ upon 10 mM Ca^2+^ addition as the HSQC spectra became highly dispersed at both pH 5.0 and pH 7.0 (Fig. 3 and Fig. S7) as found previously for RBD1^WT^ at pH 7.4 (*11*). The Ca^2+^-binding studies by NMR were confirmed via monitoring changes in tryptophan fluorescence whereby Ca^2+^-binding to RBD1^WT^ was observed (^Ca^K_D_ = 120 ± 20 μM; Fig. 4). Whereas no detectable effect on tryptophan fluorescence was observed for RBD1^D623A/D734A^ upon Ca^2+^ ion addition even at concentrations as high as 10 mM (^Ca^K_D_ > 10 mM). Under identical conditions, toxicity to *Vero* cells was collected upon addition of CDT having either full-length CDTb^WT^ or CDTb^D623A/D734A^. The optimal CDTa:CDTb ratio for CDT was maintained at a ratio of 1:7 throughout the assay for both toxin constructs as described previously (*29*). In these studies, only an ∼8-fold decrease in cytotoxicity was detected for CDTb^D623A/D734A^ (TC_50_ = 560±60 pM) versus CDTb^WT^ (TC_50_ = 70±20 pM) (Fig. S8).

**Fig. 3.**
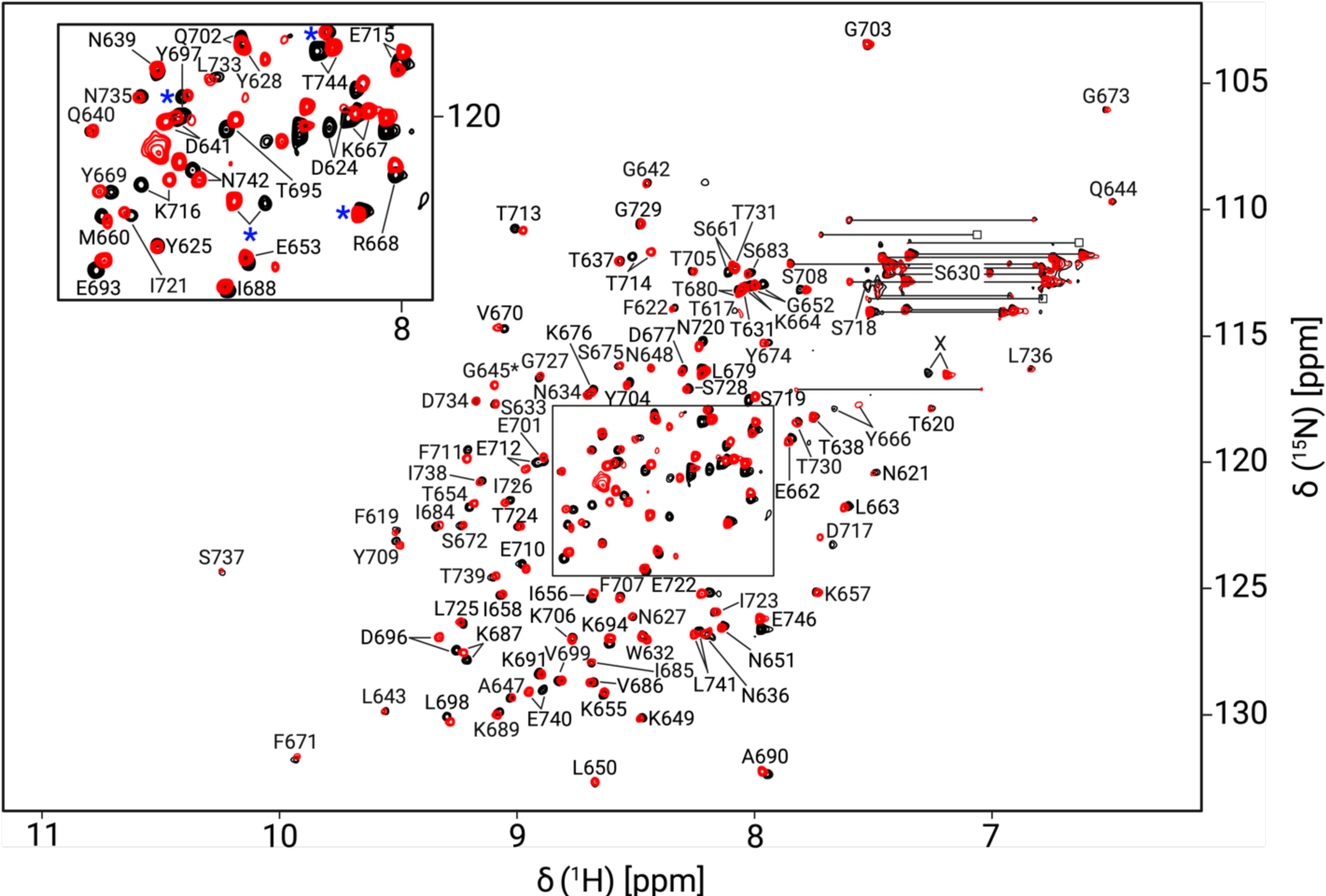
Overlay of NMR data at neutral (black) and acidic (red) pH values for CDTb (residues 616-748), termed RBD1^WT^. ^15^N-HSQC NMR data were collected at 0.1 mM RBD1^WT^, at pH 7.0 (black) and pH 5.0 (red) after 20 hours incubation with 10 mM Ca^2+^. Residues marked by blue asterisks arise from the His-tag. The contour marked “X” could not be assigned unambiguously as correlations to this H^N^ were very weak in 3D NMR data sets (not shown). The contour labeled G645* appeared in the noise at pH 7.0 but was much stronger at pH 5.0. Correlations connected by horizontal lines correspond to sidechain NH_2_ groups.

**Fig. 4.**
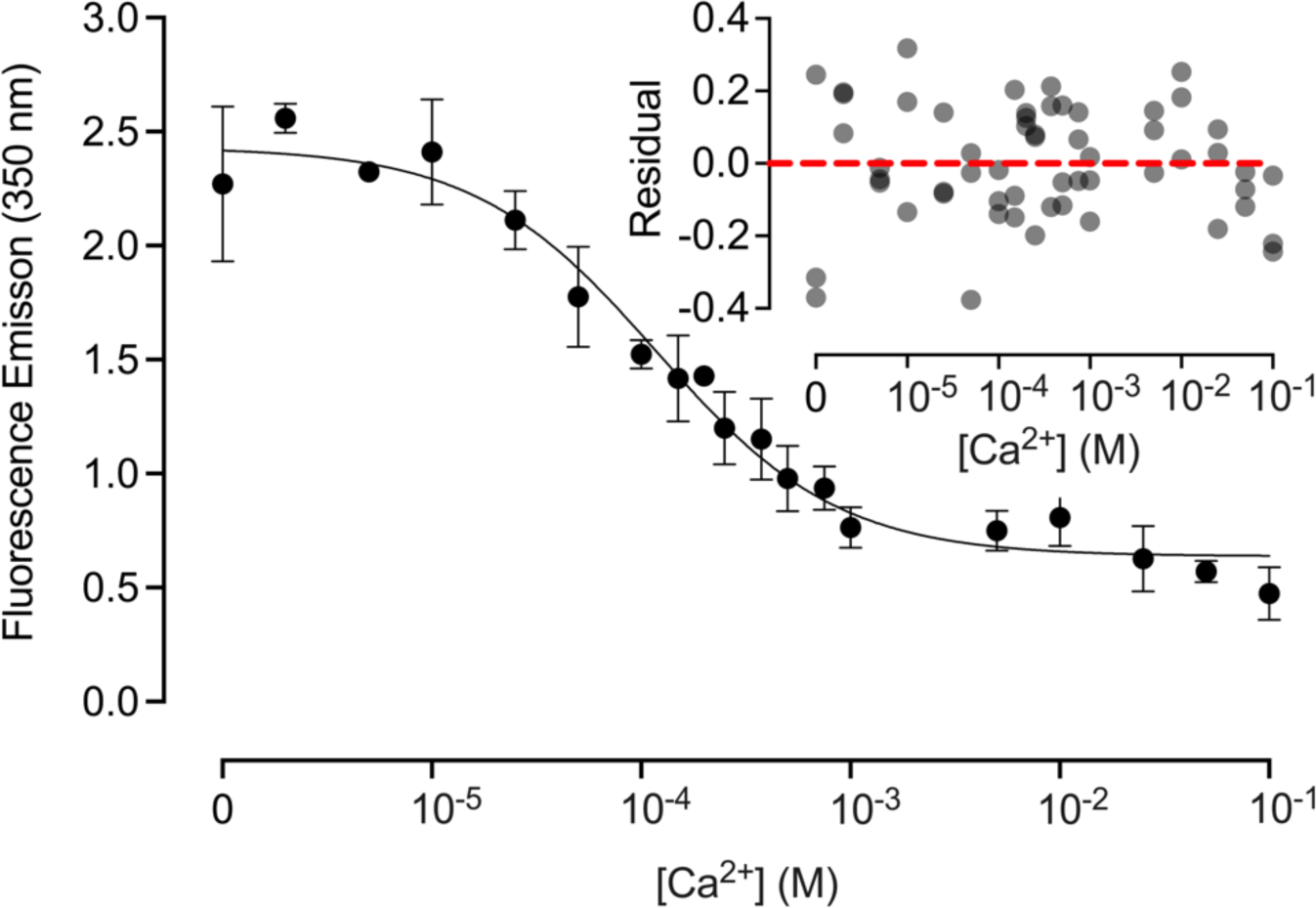
Calcium ion binding to RBD1^WT^ determined by fluorescence spectroscopy. Ca^2+^-binding to the RBD1^WT^ domain (residues 616 - 748) was monitored by changes in fluorescence emission intensity of its single tryptophan residue, Trp632. (Insert) A residual plot for the data fitting a model with noncooperative binding of Ca^2+^ to a single site in RBD1^WT^ (^Ca^K_D_ = 120 ± 20 μM).

Importantly, cytotoxicity studies showed that CDT retained a highly potent cytotoxic effect even when the Ca^2+^-binding site of RBD1 was fully ablated (Fig. 5), which is consistent with a Ca^2+^-free RBD1 domain being sufficient for delivering a highly toxic effect in the pM range.

**Fig. 5.**
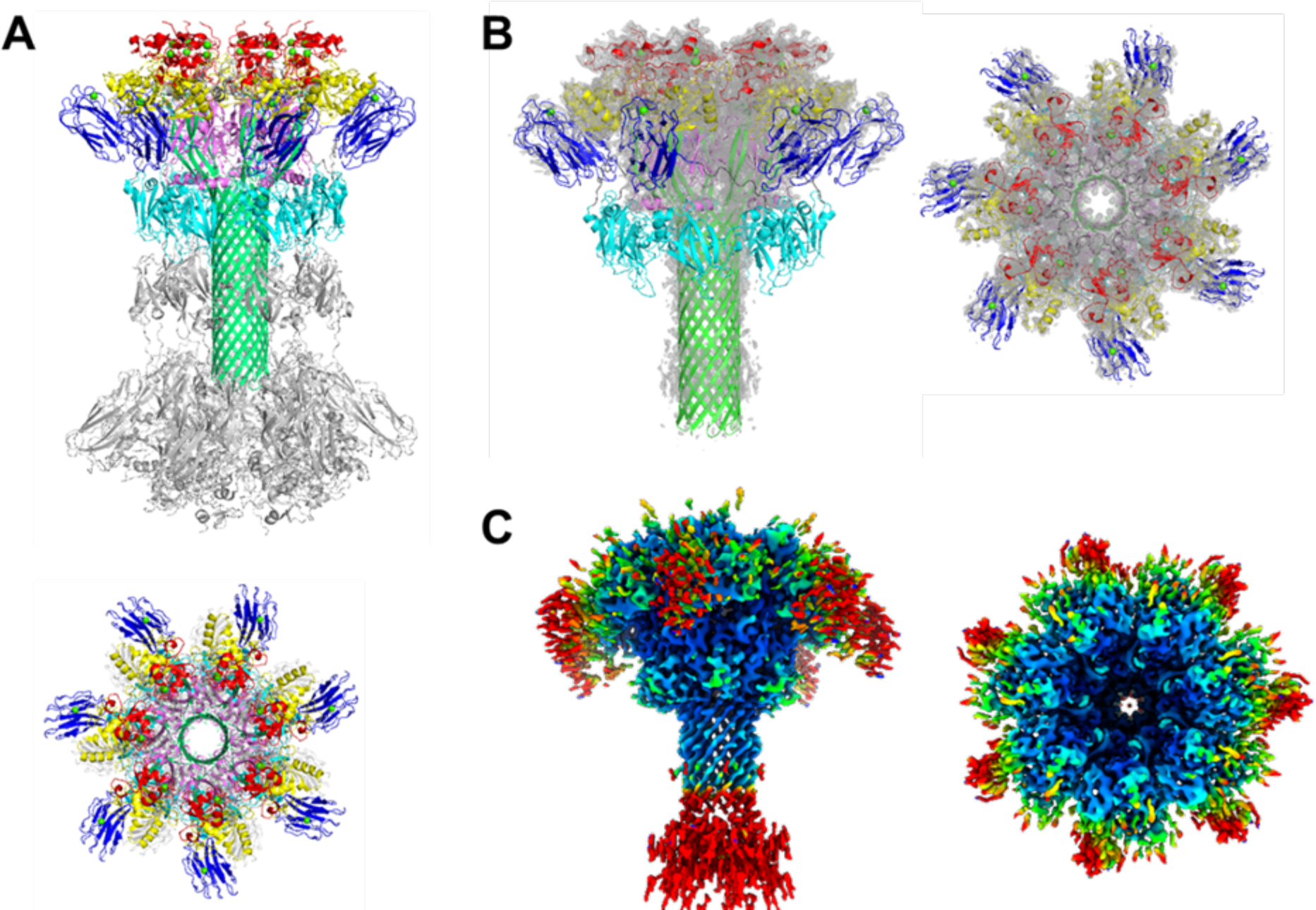
Comparison of the structural architecture of the CDTb and the CDTb^D623A/D734A^. (*A*) Side and top projections of the di-heptamer structure in the “asymmetric form” of CDTb (PDB: 6UWR). The heptamer with the β-barrel extension is illustrated in different colors; heptamerization domain 1 (HD1; residues 212-297) in red, β-barrel domain (βBD; residues 298-401) in green, heptamerization domain 2 (HD2; residues 402-486) in violet, linker region 1 (L1; residues 487-513) in grey, heptamerization domain 3 (HD3; residues 514-615) in yellow, receptor binding domain 1 (RBD1; residues 616-744) in blue, linker region 2 (L2; residues 745-756) in grey, and receptor binding domain 2 (RBD2; residues 757-876) in cyan. Dual Ca^2+^ ions are bound in HD1 and the single Ca^2+^ ion bound in RBD1 are shown as green spheres. (*B*) The single heptamer extracted from CDTb is superimposed with the electron mesh map of the CDTb^D623A/D734A^ structure. (*C*) The electron density map of CDTb^D623A/D734A^ is colored by local resolution with C7 symmetry imposed. Increased flexibility is observed in the outer regions of the core heptamer, and it is most pronounced for the RBD1 domain and the tip of the β-barrel extension.

Notably, the structure of CDTb^D623A/D734A^ was solved and found to be a single heptamer at 3.56 Å resolution (Fig. 5), much like that found for the CDTb structure solved under Ca^2+^-depleted states. Of interest is that the structure of CDTb^D623A/D734A^ lacks clear density for the RBD1 and RBD2 domains, consistent with these domains being dynamic when the RBD1 Ca^2+^ ion binding site is mutated. Interestingly, the observed heptameric structure of CDTb^D623A/D734A^ resembles a single ‘pore-forming state’ with an elongated β-barrel assembly (Fig. S9). Of further interest, our structures illustrate the existence of a single heptamer unit in soluble conditions for the first time, without detergents or lipid membrane-like support (Fig. 5C). That isolated RBD2 is monomeric in solution and crystallizes as a monomer further demonstrates that the donut-shaped RBD2 structure found in the di-heptamer oligomerization state is assisted by heptamerization and that RBD1 domains stabilize the overall protein framework (*11*). Together, these data are consistent with a mechanism by which increased flexibility of RBD1 upon Ca^2+^ removal results in destabilization of the RBD2 domain position, thus disrupting CDTb di-heptamer formation and further facilitating CDTb membrane binding and pore formation on its own. Furthermore, the ability of CDTb^WT^ alone to bind membrane could facilitate a more rapid interaction with a host cell receptor versus the RBD1 mutant, CDTb^D623A/D734A^, via membrane-lateral diffusion and a 2-versus 3-dimensional approach (i.e., for k_on_), as is the case for other beta-barrel forming bacterial toxins (*30*).

Interestingly, the β-barrel domain of CDTb (residues 298 to 401) originates from the heptameric core and elongates to form a “stem-like” structure that is essential for membrane insertion. The CDTb^D623A/D734A^ structure depicted weaker density at the end of the β-barrel and therefore residues ranging from 332 to 362 could not be modeled (Fig. 5B). This could be due to partial unfolding of the tip of the β-barrel in the absence of supporting membrane structure and/or from some of the dynamic features associated with the lack of Ca^2+^ bound to RBD1. Also intact was the structural orientation of Phe455, the key residue in “φ-gate” assembly, which in this structure agrees with the narrowest 6 Å clamp identified for the CDTb pore state (Fig. S10).

Coincidentally, the receptor binding domains (RBD1 and RBD2) were relatively disordered in CDTb^D623A/D734A^, as compared to wild-type CDTb (9) and modeling partial density in RBD1 for the double mutant could not be achieved globally. Although, when the CDTb^D623A/D734A^ map was examined with C1 symmetry, the occurrence of stronger RBD1 density was noticed in one or two of the protomers versus others (Fig. S11); whereas application of C7 symmetry averaged out density in all the RBD1 protomers among the seven subunits. Thus, the absence of a bound Ca^2+^ ion in the RBD1 domain of CDTb^D623A/D734A^ leads to a less defined structure in the RBD domains than for wild-type CDTb, and the increased flexibility of RBD1 in the CDTb^D623A/D734A^ Ca^2+^ binding site mutant was found to affect stability of the RBD2 domains and prevent observation of the assembly of RBD2 “donut” structures observed previously for the wild type di-heptamer CDTb structures (9).

### Ca^2+^ depletion upon CDTb entering early endosomes can trigger CDTb membrane binding and pore formation

From the CDTb structures presented here, we highlight that removal of the Ca^2+^ site in RBD1 by mutation or Ca^2+^ dissociation via dialysis results in a functional CDTb binary toxin component at pM levels. To understand better why low pH alone does not trigger pore formation, as observed for anthrax toxin, we compared the superimposed prepore and pore states of CDTb and anthrax PA. While the heptamerization core is well aligned in both proteins, a key difference in the β-barrel forming domain was detected in the prepore state overlay (Fig. S12). A direct link of β-barrel domain and RBD1 in CDTb restricts mobility of the β-loop and stabilizes the prepore assembly. Therefore, Ca^2+^ removal sacrifices the stability of RBD1, triggering a conformational rearrangement in the CDTb membrane insertion β-loop to transition from prepore state to pore state. Thus, the overall mechanism for RBD1 Ca^2+^-sensing is that the stable Ca^2+^ bound conformation of RBD1 forms direct interactions with the β-barrel domain, locking it into an orientation that is folded into the heptamerization domain. Whereas Ca^2+^ depletion in maturing endosomes unfolds RBD1, thus freeing the β-barrel domain to reassemble into the heptameric membrane binding and pore-forming conformation (Fig. 6).

**Fig. 6.**
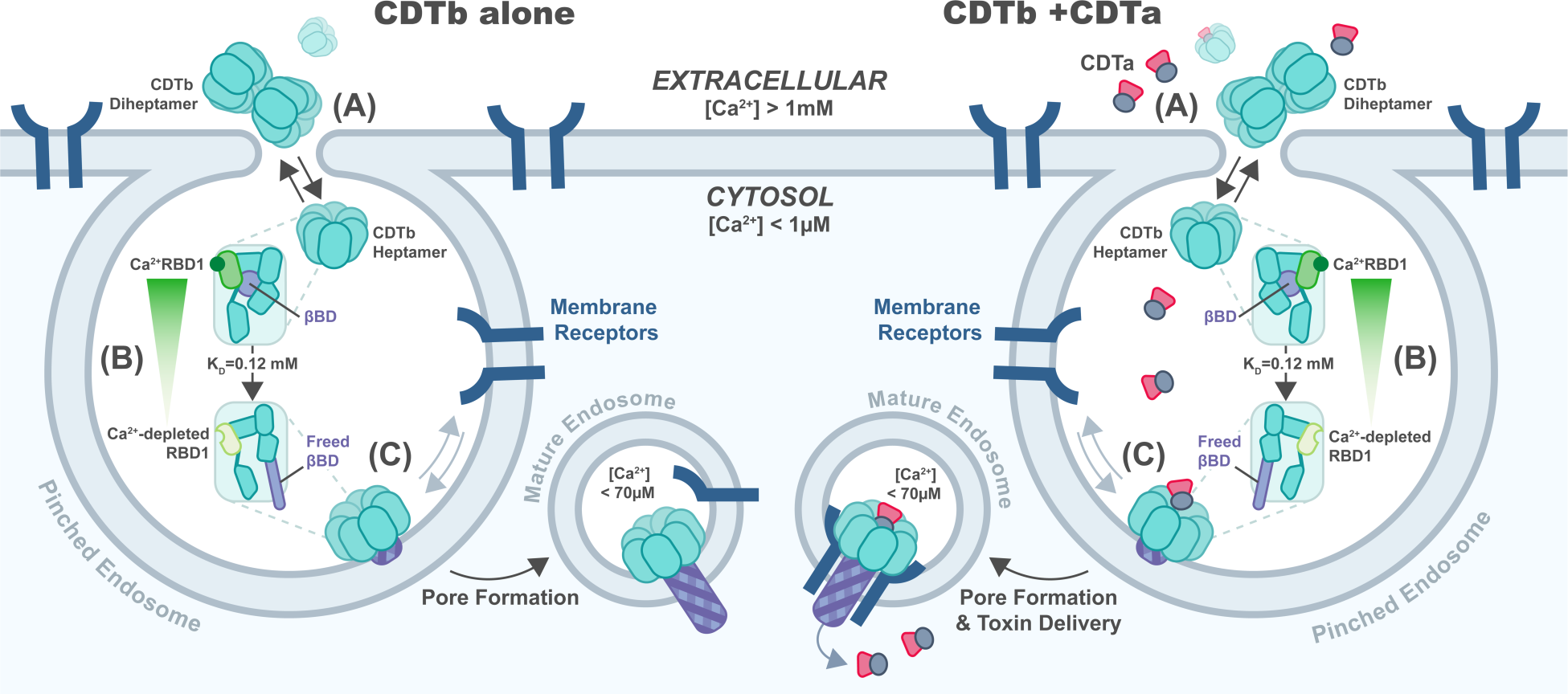
Model for membrane association and pore-formation for CDTb alone and in the presence of CDTa. In an extracellular environment ([Ca^2+^] > 2 mM), CDTb exists in equilibrium between single subunit, heptamer, and di-heptamer states (*11*), in which Ca^2+^ is bound to RBD1 and HD1, but is unable to bind lipid bilayers or form pores. Upon dissociation of Ca^2+^ from the RBD1 domain of CDTb (^Ca^K_D_ = 120 ± 20 µM), the RBD1 domain is in conformational exchange allowing the β-barrel forming residues to reassemble as needed to bind lipid bilayers and form pores. A change in orientation of beta-barrel forming residues (purple) within single subunits of the CDTb heptamer are presented as CDTb transitions from an extracellular (pH = 7.4 and [Ca^2+^] > 1 mM) to an endosomal environment (pH ∼ 5.5 and [Ca^2+^] < 70 µM). The membrane-penetrating partial β-barrel (pre-insertion state) formation is initiated upon endocytosis. Low internal Ca^2+^ concentration (< 70 µM) and acidic pH (∼ 5.5) in the endosome ensure the membrane-spanning pore formation with the complete extended β-barrel. (Right) Components needed for a “toxic” outcome with CDTa and CDTb present under cellular conditions are also shown on the right. In the presence or absence of CDTa, CDTb-receptor interactions could be facilitated via membrane lateral diffusion of CDTb and/or receptor(s); however, a complete mechanism of action for CDTa delivery by CDTb within a mature endosome requires further examination.

In summary, the data presented here are consistent with a novel mechanism of host cell engagement by the cell-binding and toxin delivery component of the binary toxin, CDTb. Specifically, Ca^2+^-dissociation from the RBD1 domain of CDTb or mutation of this site were found to be sufficient for this component of CDT to bind and pierce lipid bilayers. Such a mechanism of action can be understood at atomic detail as it is clear by NMR and cryoEM studies that dissociation of Ca^2+^ from this RBD1 site induces conformational dynamics that directly impact a di-heptamer to heptamer transition, membrane binding, and pore formation. Physiologically, these findings are consistent with a molecular mechanism in which entry of CDTb into early endosomes is sufficient for Ca^2+^-dissociation from RBD1 followed by CDTb membrane binding and pore formation that pierces membrane separating the endosome from the cytosol. The absence of Ca^2+^ bound to the RBD1 site also provides a heptamer state of CDTb that is highly dynamic, which is a physical attribute likely to be important for these molecular events. This study now provides three mechanistic possibilities that can be used by bacterial translocases of this class to trigger timely activation from within maturing endosomes including (i) lowering pH (i.e., anthrax), (ii) lowering free Ca^2+^ ion concentration (i.e., CDT), and/or (iii) a combination of both physiologically relevant mechanisms. For the *C. difficile* binary toxin, how host cell receptor and/or CDTa binding to CDTb regulates CDT entry into host cells requires further study.

## Acknowledgments

The National Institute of Standards and Technology (NIST) Center for Nanoscale Science and Technology cleanroom in Gaithersburg, MD is acknowledged for the preparation of chips needed for SPR/EIS experiments. We express our appreciation to John J. Kasianowicz for sharing the SPR/EIS data collected for anthrax toxin.

## Funding

National Institutes of Health grant R56AI152397 (DJW, AdG) National Institutes of Health grant R01GM129327 (DJW) Center for Biomolecular Therapeutics (DJW) Maryland Center for Advanced Molecular Analyses (DJW, EP) University of Maryland - Institute for Bioscience and Biotechnology Research (IBBR) University of Maryland School of Medicine Maryland Department of Health’s Restitution Fund Program (CH-649-CRF) and NCI-CCSG (P30CA134274)

## Author contributions

Conceptualization: EP, AdG, DJW Methodology: DLA, SES, KAA, RG-R, KMV, IFN, AXS, RRR, VIS, EP, DJW Investigation: DLA, SES, KAA, RG-R, KMV, IFN, AXS, RRR, VIS, AdG, EP, DJW Funding acquisition: EP, AdG, DJW Supervision: EP, KMV, VIS, DJW Writing – original draft: DLA, KAA, RG-R, KMV, SES, RRR, VIS, MEC, EP, DJW Writing – review & editing: DLA, KAA, RG-R, KMV, SES, RRR, VIS, AdG, MEC, EP, DJW

## Competing interests

The authors declare no competing interest.

## Data and materials availability

Cryoelectron microscopy density maps have been deposited in the Electron Microscopy Data Bank (EMDB), https://www.ebi.ac.uk/pdbe/emdb/ (accession nos. EMD-28207 for CDTb^D623A/D734A^, EMD-28206 for CDTb^(-Ca)^ class 1 and EMD-28205 for CDTb^(-Ca)^ class 2). Model coordinates have been deposited in the Protein Data Bank (PDB), https://www.wwpdb.org/ (PDB ID codes 8EKM for CDTb^D623A/D734A^, 8EKL for CDTb^(-Ca)^ class 1, and 8EKK for CDTb^(-Ca)^ class 2).

All other data are available from the corresponding authors upon request.

## Supplementary Materials

Materials and Methods Figs. S1 to S12

Table S1 References (*31–40*)

## Materials and Methods

### Protein Expression and Purification

Active CDTa^C2A^, CDTb^WT^ and the CDTb^RBD1^ domain of CDTb (residues 616-748) were expressed and purified as previously described (6, 11). To remove Ca^2+^ from CDTb^RBD1^ and CDTb^WT^, the purified proteins were dialyzed into 15 mM HEPES buffer, pH 7.0, 150 mM NaCl, 2 mM ethylenediaminetetraacetic acid (EDTA), and 2 mM triethylene glycol diamine tetraacetic acid (EGTA) at 4 °C overnight. A second dialysis into 15 mM HEPES buffer, pH 7.0, and 150 mM NaCl was performed overnight at 4°C, to remove EDTA and EGTA. To test whether any residual EDTA affected membrane binding and pore formation, a fraction of CDTb^WT^ was passed through a Sephedex G25 column equilibrated 20 mM HEPES buffer, pH 7.0, and 50 mM NaCl to eliminate any residual EDTA. No difference was observed when membrane binding and pore formation for CDTb^WT^ were compared before and after Sephadex G25 size exclusion chromatography (SEC).

The Ca^2+^-binding RBD1 mutant proteins (CDTb^D623A/D734A^, RBD1^D623A/D734A^) were engineered into pET21a (Amp^R^) DNA expression plasmids, codon optimized by TOPGene Technologies, and engineered with an N-terminal 6xHis-tag for purification. Purification of the mutant constructs followed procedures analogous to those of wild-type CDTb^WT^ and CTBb^RBD1^ (residues 616-748), respectively (6,11), with some modifications. Briefly, proCDTb^D623A/D734A^ expression was induced in by 0.5 mM isopropyl β-D-1-thiogalactopyranoside (IPTG) addition once cell cultures reached an OD_600_ of 0.6, at which point they were incubated at 37 °C for an additional 4 hours. Cells were harvested by 15 minutes centrifugation at 5000 rpm, at 4 °C, and the cell pellet was resuspended in lysis buffer containing 20 mM Tris at pH 8.0, 300 mM NaCl, 5 mM β-mercaptoethanol (βME), and 200 mM phenylmethylsulfonyl fluoride (PMSF) to inhibit cellular proteases. Cell lysis involved 3 cycles of sonication while cooled, and cell debris was removed by centrifugation at 15000 rpm, for 45 min at 4 °C. A filtered clear supernatant was loaded onto a HiPrep 16/60 IMAC column and purified (>95%). Pure fractions of proCDTb^D623A/D734A^ were pooled and dialyzed into 15 mM HEPES buffer, pH 7.0, 150 mM NaCl, and 10% glycerol prior to activation. Active CDTb^D623A/D734A^ was obtained by proteolytic cleavage of the N-terminal activation domain with a slight modification from that used for wild-type CDTb (6,11). Specifically, for active CDTb^D623A/D734A^ preparations, the cleavage reaction was achieved by overnight incubation with chymotrypsin at 20 °C. After >90% of cleavage was confirmed by SDS-PAGE, activated CDTb^D623A/D734A^ protein was injected onto Superdex S200 size exclusion column equilibrated 15 mM HEPES, pH 7.0, 150 mM NaCl, and 0.5 mM dithiothreitol (DTT) with the purified protein elution (∼70 kDa; >95%) being pooled, concentrated, and stored at −80 °C.

Wild-type or Ca^2+^-mutant plasmids of RBD1 (residues 616-748), termed RBD1^WT^ and RBD1^D623A/D734A^, respectively, were engineered. Protein production occurred in 2 L of [^15^N]-labeled MOPS media or 2 L LB media for ^15^N-labeled and unlabeled expression, respectively. Overexpression was induced with 0.5 mM IPTG when cultures reached OD_600_ values of approximately 0.6, followed by continued induction at 25 °C for 18 hours. Cells were harvested by centrifugation at 4000 rpm for 30 minutes at 4 °C, resuspended in buffer containing 50 mM Tris-HCl, pH 8.0, 500 mM NaCl, and 10 mM βME, and lysed as described (6, 11). Purification was achieved via Ni Sepharose™ 6 Fast Flow or Ni NTA-agarose columns and fractions containing RBD1^WT^ or RBD1^D623A/D734A^ (∼16.4 kDa) were pooled and dialyzed into 15 mM HEPES, pH 7.0, 150 mM NaCl, and 0.5 mM TCEP before being concentrated and stored at −80 °C.

### Cryo-EM Sample Preparation, Data Acquisition, and Processing

A 3 µL aliquot of purified CDTb^D623A/D734A^protein sample at a concentration of 30 µM was applied to a glow discharged QUANTIFOIL® copper R 1.2/1.3 grid with a 2 nm thin extra carbon layer (QUANTIFOIL, GMBH). For calcium depleted CDTb (28 µM), glow discharged QUANTIFOIL® copper R 2/2 grid with a 2 nm thin extra carbon layer (QUANTIFOIL, GMBH) was used. Grids were double blotted for 2.5 seconds using a FEI Vitrobot IV with a waiting time of 0 s, in 100% humidity. The sample grids were plunged into liquid ethane for vitrification. Frozen grids were stored in liquid nitrogen until data collection.

Motion corrected micrographs were produced, and contrast transfer function estimates were obtained in CryoSPARC (*31, 32*). For the CDTb^D623A/D734A^, a total of 993,305 particle images were picked by template picker, and several rounds of 2D classifications were performed in cryoSPARC to select the most resolved particles. A total of 126,516 particles with different orientations were re-extracted to build the initial model of CDTb^D623A/D734A^. The selected particles were used for *ab initio* model generation with 15 classes to clean out the particle stack. The classes with distinguishable initial volumes were selected for further 3D refinement and CTF refinement. Final 3D Homogeneous refinement of 78,587 particles at C1 symmetry yielded a high-resolution density map at 4.04 Å. C7 symmetry refinement of the structure resulted in a density map with an improved resolution at 3.56 Å. Resolutions were estimated based on the Fourier Shell Correlation (FSC).

For Ca^2+^ depleted CDTb, a total of 1,109,860 particles were picked by template picker, and multiple rounds of 2D classifications were performed in cryoSPARC to screen the particles with high resolution. A total of 105,207 particles with different orientations were re-extracted to generate *ab-initio* models. Out of the 10 *ab-initio* models generated, 8 classes consisting of 91,077 particles were selected for another round of *ab-initio* modeling. The classes generated revealed two distinguishable initial models including a ‘flower-on-a-stem’ form with and without the RBD2 ring at the bottom. The volumes with and without the RBD2 electron density consisting of 36,395 and 33,531 particles, respectively, were used for 3D refinement separately. Non-uniform refinement in cryoSPARC using the two initial models yielded two high resolution density maps, at 3.71 Å and 3.61 Å resolution. C7 symmetry refinements of the two structures were carried out in cryoSPARC and yielded density maps for both heptamers with and without the RBD2 ring with improved resolutions, at 3.28 Å and 3.06 Å resolution, respectively. Resolutions were estimated based on the Fourier Shell Correlation (FSC) with 0.143 as cutoff (Table S1).

### Model Building and Refinement

To build the models for CDTb^D623A/D734A^ and CDTb^(-Ca)^, a single heptamer assembly of the asymmetric CDTb structure (PDB: 6UWR) with the β-barrel was used as a starting model and fitted into the electron density maps of CDTb variants using UCSF Chimera (*33*). Then, the models were manually adjusted in COOT and refined using real space refinement in PHENIX (*34, 35*).

### Protein-Lipid Interaction by SPR/EIS

To prepare the tethered bilayer lipid membrane (tBLM) on the gold-coated sapphire chip, a thiol lipid mimic 1,2-di-O-myristyl-3-[x-mercaptohexa(ethylene oxide) glycerol] (WC14) was used (synthesized by Dr. David Vanderah) as the tethering molecule (*36*). Distilled 2-Mercaptoethanol (βME) (purity > 99%) was used as a space-filling component between WC14 molecules. Powder of the lipid (purity > 99%) 1-palmitoyl-2-oleoyl-snglycero-3-phosphocholine (POPC, Avanti Polar Lipids) was dissolved in anhydrous ethanol (purity > 99.5%), to a final concentration of 10 mM. The gold metal deposition on a sapphire chip for the SPR-EIS experiment was performed by magnetron sputtering on a Denton Vacuum Discovery 550 Sputtering System at the NIST Center for Nanoscale Science and Technology cleanroom in Gaithersburg, MD, USA. Substrate was then incubated overnight in a 0.2 mM ethanol solution of WC14/βME with a molar ratio of 25:75 to form a mixed self-assembled monolayer (SAM) of WC14/βME. After rinsing with ethanol and drying in a nitrogen stream, the SAM-coated chip was installed onto the home-built SPR/EIS setup with the cylindrical SPR/EIS cell (1 mL volume) equipped with a perfluoro elastomer Kalrez O-ring (i.d. = 6 mm) against Au/SAM surface. To ensure optical contact between sapphire prism and sapphire chip a few drops of 1-iodonaphthalene (ƞ = 1.70) was used when mounting the chip to the instrument. A tBLM was formed with the fast solvent exchange method (*37*). For the tBLM, 40 µL of a phospholipid (PL) solution was transferred into the SPR/EIS cell with the SAM-coated chip. After 1 min of incubation, 20 µL of PL solution was removed carefully and after a total of 2 min incubation, it was rigorously rinsed with 1 mL of 20 mM HEPES buffer at pH 7.0, and 50 mM NaCl for 10 repetitive cycles. Formation of PL molecules was confirmed by EIS measurements. The PL overlayers was eliminated from the tBLM by incubation in a 10% to 20% ethanol solution in water for 10 min. Ethanol solution was replaced with water in the EIS/SPR cell followed by rinse with 950 µL of buffer (during washing cycles 50 µL was left to in the cell to avoid the tBLM contact with air). Buffer replacement was repeated for 7-8 times and left undisturbed for ∼1 hour to ensure a homogenous stable bilayer on the tBLM before protein introduction. Then, pH 7 buffer was replaced by 20 mM NaOAc buffer at pH 5.5 and 50 mM NaCl and left undisturbed for ∼40 minutes to ensure a homogenous stable bilayer on the tBLM before protein introduction. Initial formation of a homogenous stable tBLM was ensured at the physiological pH of 7.0. Buffer exchange to low pH buffer increases the electrical resistance of the lipid bilayer because of the protonation and the decrease in charge of the phospholipid phosphate groups. Hence, the electrical conductance decreases. In addition, low pH results in a decrease in SPR signal of phospholipid bilayer. After successful formulation and stability of a tBLM was confirmed by EIS measurements, 1 µM of protein was introduced. Protein adsorption was monitored by SPR and EIS measurements. Successful formulation and stability of a tBLM was confirmed by EIS measurements. The characteristic small semicircle in the EIS spectrum confirmed the bilayer association. EIS data fitting was conducted using the ZView software (Scribner Associates, Inc.) to derive capacitance and resistivity values of the bilayer behavior. While change in SPR signal was recorded every 2 s for the duration of the experiment, EIS measurements were recorded every 5 min intervals for >1 hour and resistance were obtained by fitting to equivalent circuit model. Adsorption of the CDTb^(-Ca)^ to the phospholipid membrane at pH 5.5 generates an influx of ions due to pore insertion. Difference in SPR reading at the beginning of protein adsorption and at the end of protein desorption provides the net change. A positive net change signified channel formation in the lipid bilayer. The same procedure was repeated with the CDTb alone. Before each new experiment, the SPR/EIS cell with the O-ring was rinsed with detergent followed by 20-30 mL of miliQ water. Then it was sonicated in ethanol and dried under a stream of nitrogen. The experiments were repeated three times to ensure reproducibility. All measurements were conducted at 25.0 ± 0.3 °C.

### NMR Spectroscopy

2D [^1^H, ^15^N]-HSQC (heteronuclear single quantum coherence) NMR experiments of 0.1 mM RBD1^WT^ in 15 mM HEPES, 150 mM NaCl, 10 mM CaCl_2_, 0.5 mM TCEP, 0.2 mM TPEN, and 20% D_2_O, from pH 7.0 to pH 5.0 at 25 °C were performed on an 800 MHz Bruker Avance NMR spectrometer. A 2D [^1^H, ^15^N]-HSQC NMR experiment of 0.25 mM RBD1^WT^ in 15 mM HEPES (pH 7.0), 150 mM NaCl, 0.5 mM TCEP, 0.2 mM TPEN, and 20% D_2_O at 25 °C was performed on an 800 MHz Bruker Avance NMR spectrometer. 10 mM CaCl_2_ was then added and 2D [^1^H, ^15^N]-HSQCs were collected under the same conditions over a 20-hour period. 2D [^1^H, ^15^N]-HSQC NMR experiments of 0.1 mM RBD1^D623A/D734A^ in 15 mM HEPES (at pH 7.0 and pH 5.0), 150 mM NaCl, 0.5 mM TCEP, 0.2 mM TPEN, and 20% D_2_O at 25 °C were performed on a 600 MHz Bruker Avance III NMR spectrometer. 10 mM CaCl_2_ was then added and 2D [^1^H, ^15^N]-HSQCs were collected under the same conditions over a 20-hour period. All NMR experiments were processed through NMRPipe and analyzed using CcpNmr Analysis (*38, 39*).

### Ca^2+^-binding as monitored by Fluorescence Spectroscopy

Ca^2+^ titrations of RBD1^WT^ and RBD1^D623A/D734A^ were carried out using a Varian Cary Eclipse fluorometer at 25 °C to monitor RBD1^WT^ Ca^2+^-binding (*40*). Samples contained 25 μM RBD1^WT^ or 40 μM RBD1^D623A/D734A^, 15 mM HEPES (pH 7.0), 150 mM NaCl, 0.5 mM TCEP, and increasing concentrations of CaCl_2_ (0 µM, 2 µM, 5 µM, 10 µM, 25 µM, 50 µM, 100 µM, 150 µM, 200 µM, 375 µM, 500 µM, 750 µM, 1 mM, 5 mM, 10 mM, 25 mM, 50 mM, 100 mM, and 500 mM (for only RBD1^D623A/D734A^), respectively. Samples were left to incubate overnight to allow to achieve equilibrium. Ca^2+^-binding was monitored via change in tryptophan emission (W632) at 350 nm upon excitation at 295 nm. The slit width for excitation and emission was 5 nm. Fluorescence Ca^2+^-binding experiments were done in triplicate and data were fit to a non-linear regression curve for log_10_([Ca^2+^]) vs. fluorescence emission intensity via a noncooperative binding model with a single binding site using Graphpad Prism 10. The results presented are average values of the three replicates with the standard error of the averages represented with error bars.

### Vero Cell Toxicity Assays of CDT

Vero cells (ATCC-CCL-81) obtained from American Type Culture Collection, were cultured in complete media EMEM (Corning Cellgro 10-009-CV) containing 10% FBS. A cell density of 4000 cells/well was maintained in 200 µL complete media placed in a 96-well plate (Corning #3904) by overnight incubation in a humidifier with 5% CO_2_ at 37 °C. To formulate the binary toxin complex, purified CDTa was combined with active CDTb^D623A/D734A^ or CDTb in complete media at the optimal molar ratio of 1:7, CDTa:CDTb. A final volume of 100 µL from each dilution was added into the each well in 96-well plate with existing media to test at toxin concentrations up to 1nM (1nM CDTa: 7nM CDTb). Vero cells were incubated with binary toxin with 5% CO_2_ at 37 °C and a dose-response curve was generated for each combination. After 24 hours, plates were centrifuged at 1000 rpm for 1 min. The supernatant was removed, and cells were fixed with 1x PBS with 3.7% formaldehyde for 15 min at room temperature. Cells were washed in 1x PBS and then incubated in 1x PBS with 1% Triton-X-100 for 5 min at room temperature, to permeabilize the cells. Then, cells were again washed with 1x PBS and incubated in the dark with 0.016 μM Alexa Fluor 488 Phalloidin in 1x PBS and 1% BSA for 1 hour at room temperature. For imaging, cells were washed again and incubated with 1x PBS. Fluorescence emission of Alexa Fluor 488 Phalloidin at 520 nm was recorded in well-scanning mode using the BMG PHERAstar FS with an excitation wavelength of 495 nm. Data were plotted using GraphPad Prism 9 and a TC_50_ was calculated using a four-parameter logistic regression analysis.

---

**Fig. S1.**
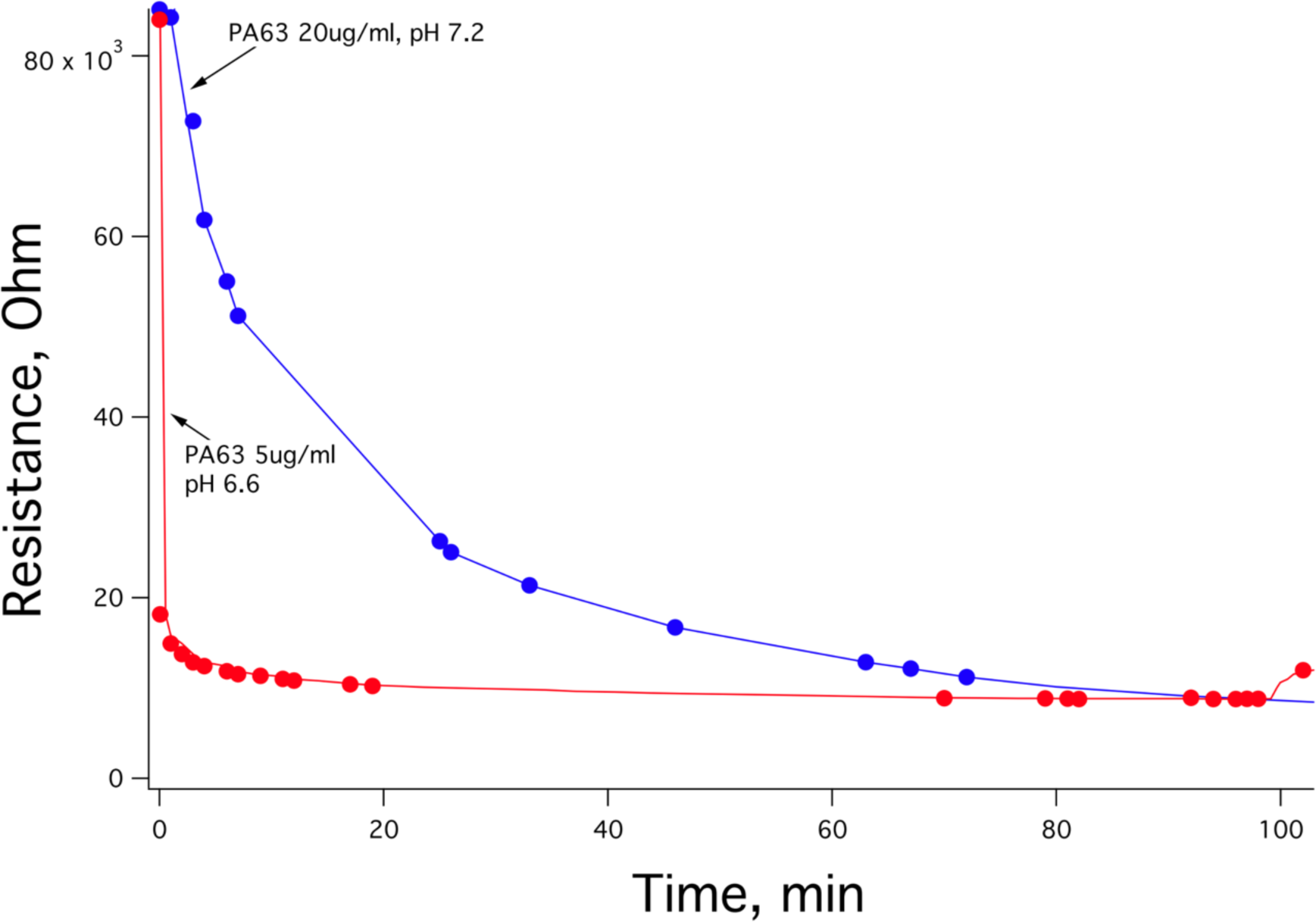
Membrane interactions of anthrax PA63 at pH 7.2 and pH 6.6. Resistance measured by electrochemical impedance spectroscopy shows rapid protein insertion into lipid membrane at lower pH (*23*).

**Fig. S2.**
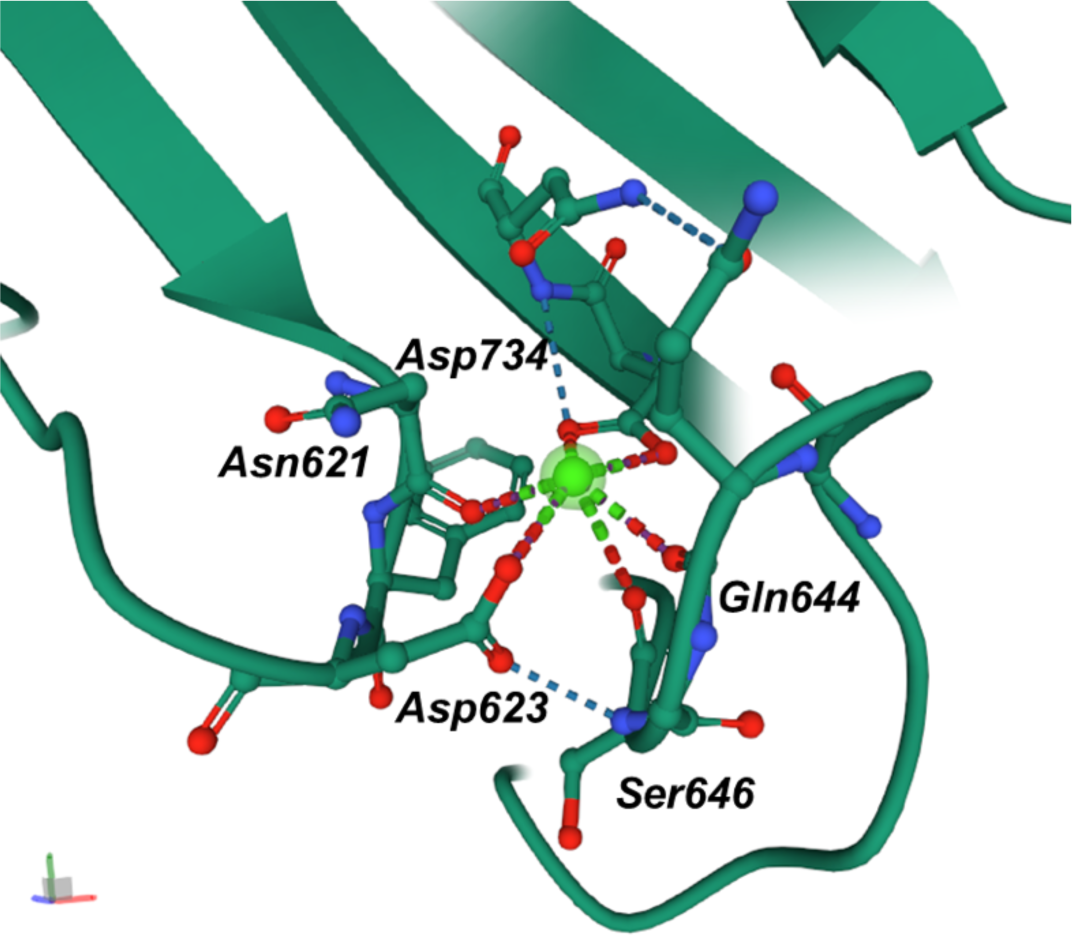
Ca^2+^ binding site in RBD1. CDTb wild type (PDB: 6UWR) contains a single Ca^2+^ ion (shown in bright green) in the RBD1 domain and the liganding oxygen atoms are from Asn621, Asp623, Ser646, Gln644, and Asp734. Point mutations were targeted at the two aspartic acid residues, Asp623 and Asp734, which were converted into alanine residues as discussed.

**Fig. S3.**
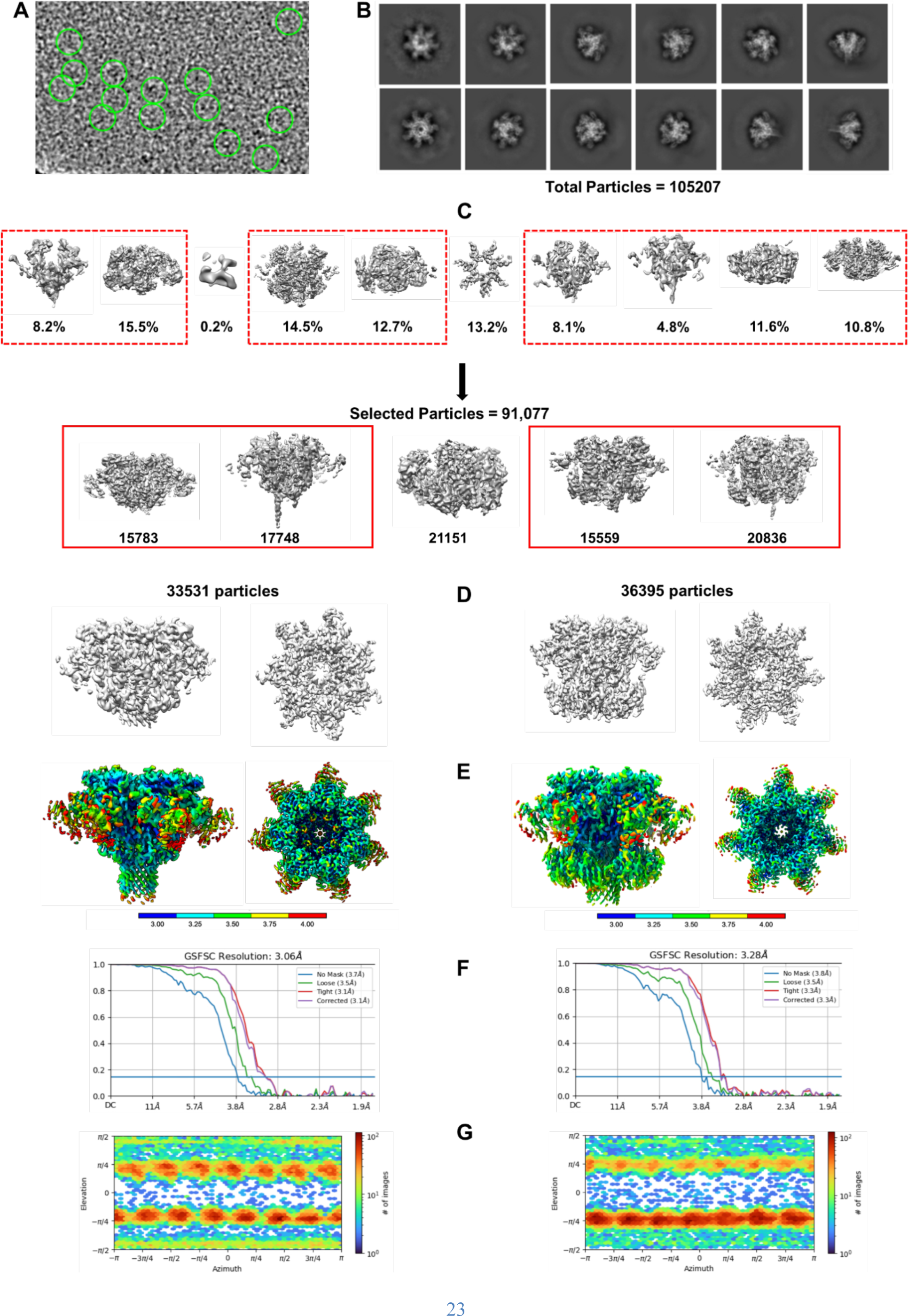
Structural characterization of Ca^2+^ depleted WT CDTb. (*A*) Example of a motion corrected micrograph with a few particles highlighted in circles to show different orientations. (*B*) Results of 2D classification generated by cryoSPARC. (*C*) Results of 3D classification generated by cryoSPARC. Classes that were combined in the final reconstructions are highlighted in red boxes. RBD2 distorted structure (class 1) is shown on the left side. The RBD2 intact structure (class 2) is shown on the right side. (*D*) Resulting electron density maps at C1 symmetry. (*E*) Resulting electron density maps colored by local resolution after application of C7 symmetry element. (*F*) FSC curves from the final reconstruction with C7 symmetry imposed. (*G*) Directional distributions of particle orientation.

**Fig. S4.**
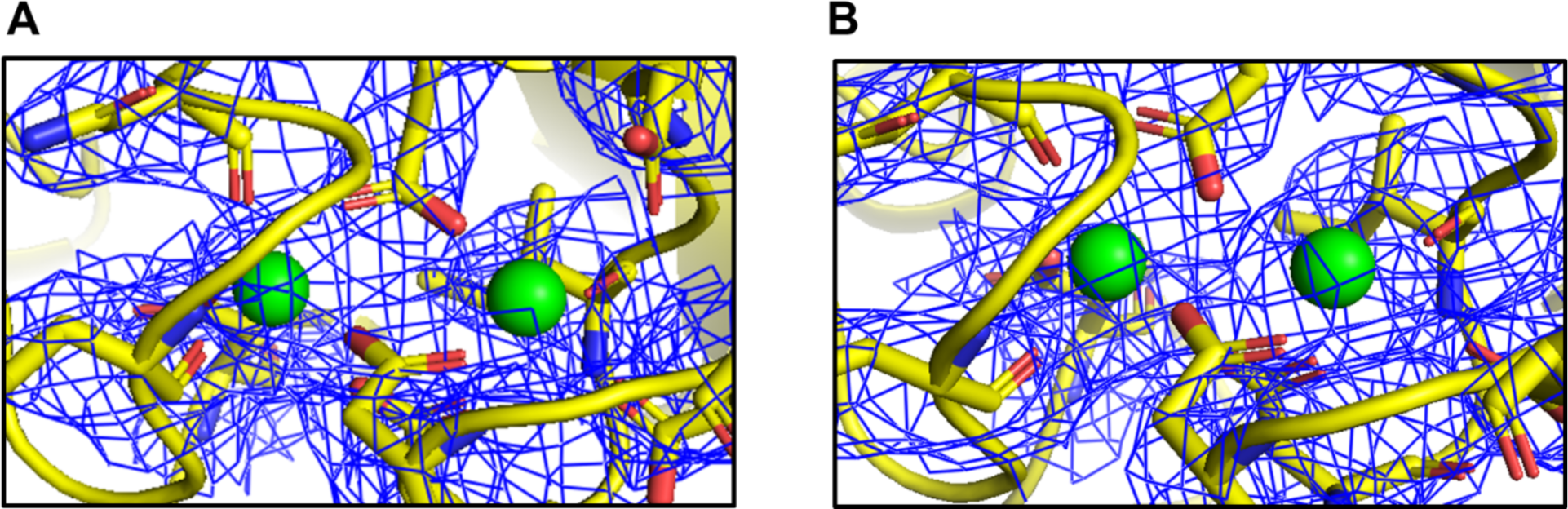
High affinity dual Ca^2+^ binding sites located in the N-terminal heptamerization domain 1 (HD1) of Ca^2+^-depleted WT CDTb. Dialysis of WT CDTb against the chelex-treated 15 mM HEPES buffer (pH 7.0), 150 mM NaCl, 2 mM EDTA, and 2 mM EGTA only removed the Ca^2+^ in RBD1 while tightly bound Ca^2+^ in HD1 remained intact. The coulomb potential maps (i.e., cryoEM electron isomesh maps; blue) in both classes of Ca^2+^-depleted WT CDTb (*A*) class 1 with increased flexibility in both RBD1 and RBD2; (*B*) class 2 with increased flexibility in RBD1 only, depicted electron density for two Ca^2+^ ions (Ca1, Ca2; green spheres) in the HD1. Calcium liganding residues identified involved oxygen atoms of D222/D224/E231/D273/N260/E263 for the Ca1 site, and D220/D222/D224/E321/D228/I226(C = O) for the Ca2 site.

**Fig. S5.**
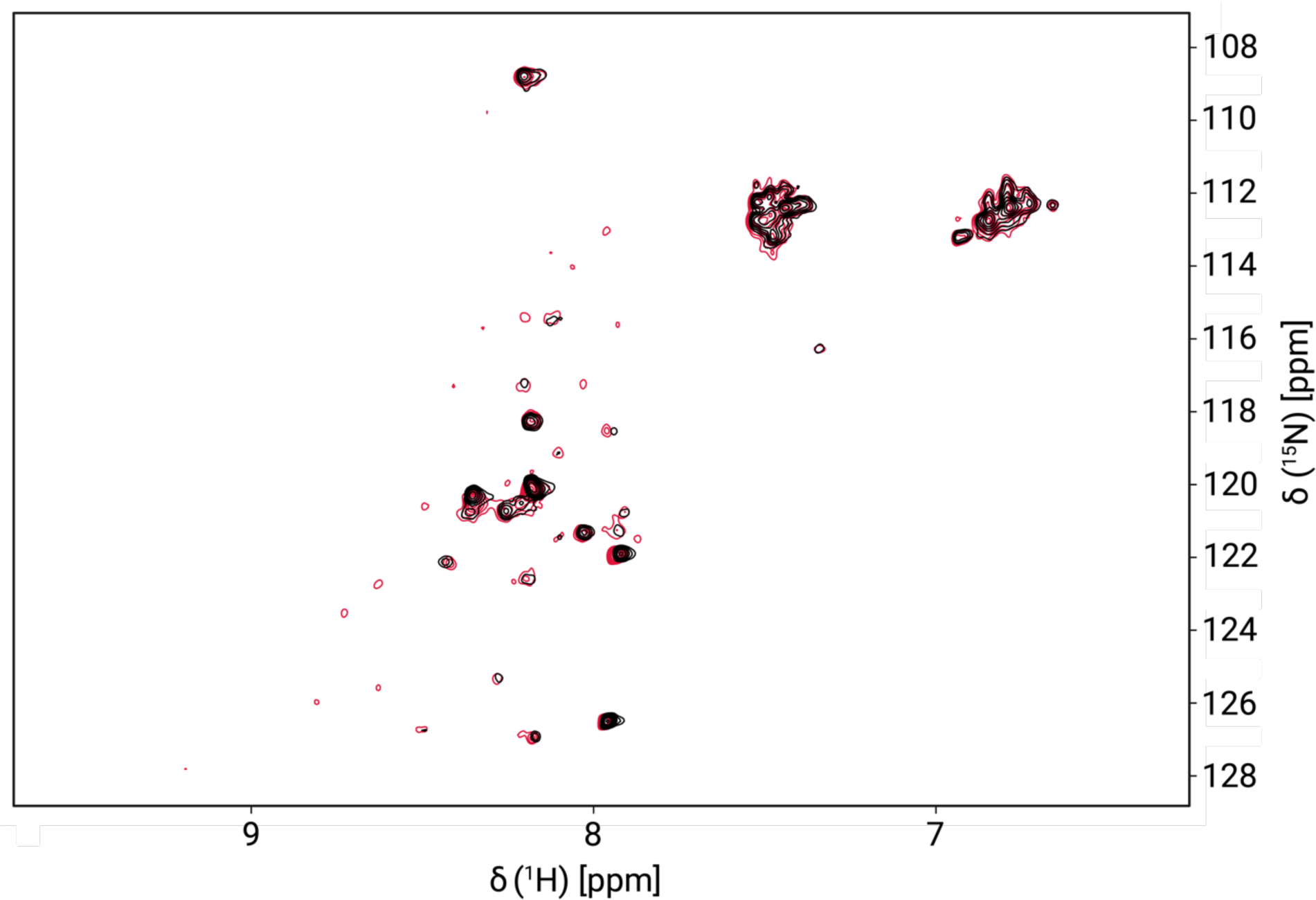
Overlay of 2D ^15^N-edited HSQC NMR spectra of RBD1^D623A/D734A^ without Ca^2+^ (black) and 20 hours after incubation with 10 mM Ca^2+^ (red). RBD1^D623A/D734A^ NMR data were collected with 0.1 mM RBD1^D623A/D734A^, 25 °C and pH 7.0.

**Fig. S6.**
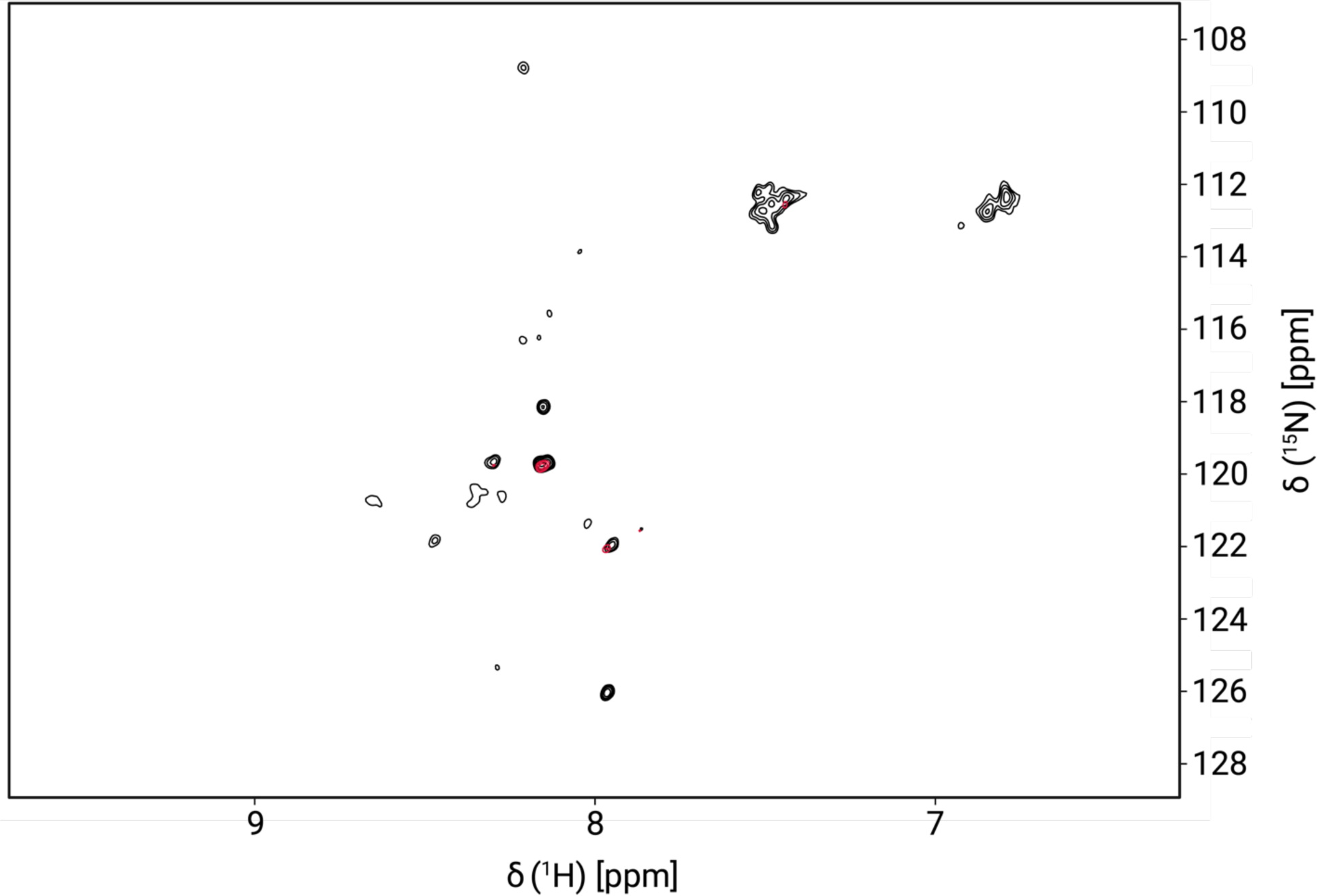
Overlay of 2D ^15^N-edited HSQC NMR spectra of RBD1^D623A/D734A^ without Ca^2+^ (black) and 20 hours after incubation with 10 mM Ca^2+^ (red). RBD1^D623A/D734A^ NMR data were collected with 0.1 mM RBD1^D623A/D734A^, 25 °C, and pH 5.0.

**Fig. S7.**
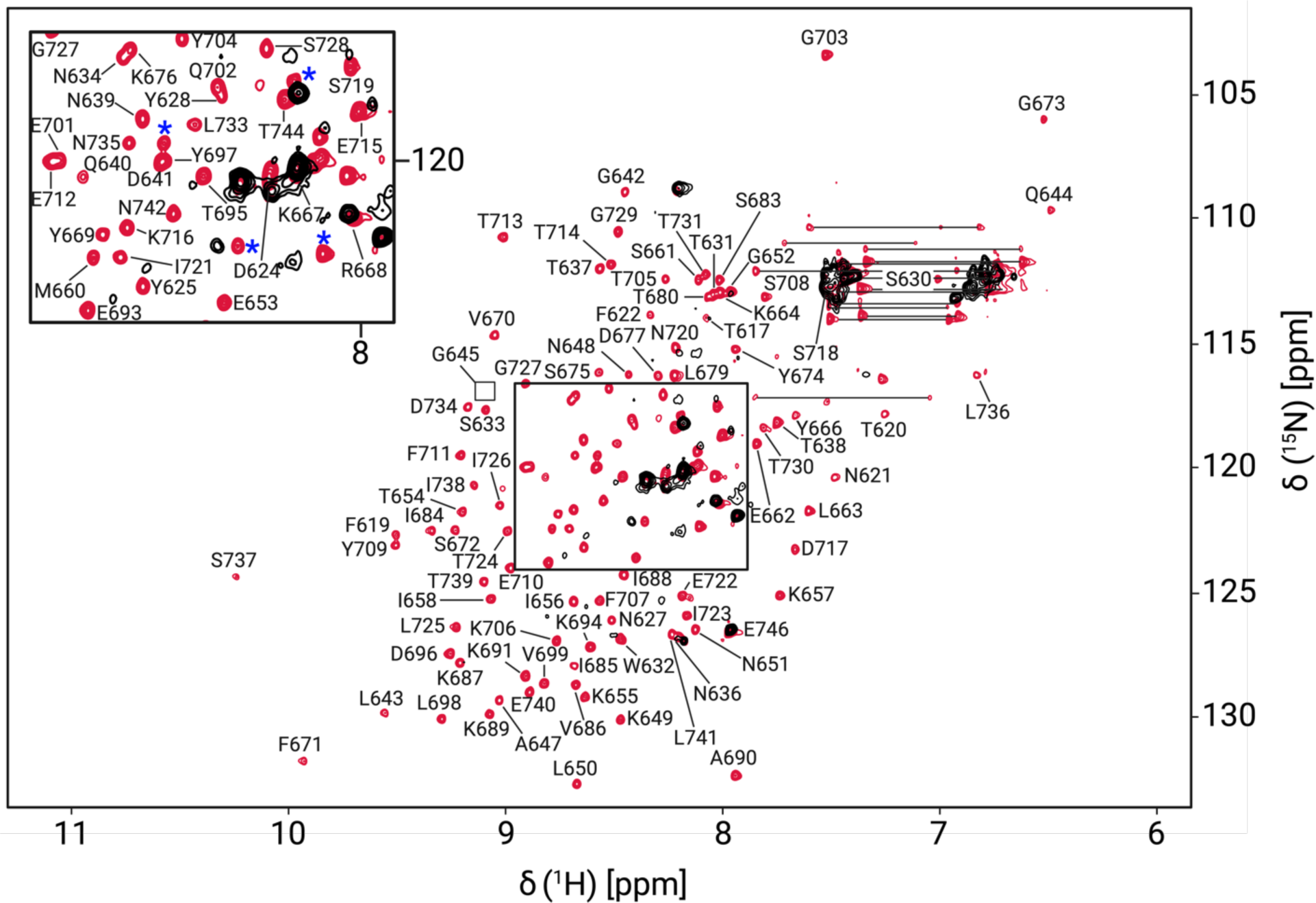
Overlay of 2D ^15^N-edted HSQC NMR spectra of RBD1^D623A/D734A^ (black) and RBD1^WT^ (red) after 20 hours incubation with 10 mM Ca^2+^. Residues marked by blue asterisks are associated with His-tag. A correlation for G645 appeared in the noise and is indicated by a box. Correlations connected by horizontal lines correspond to sidechain NH_2_ groups. RBD1^WT^ data were collected with 0.25 mM RBD1^WT^, 25 °C and pH 7.0; RBD1^D623A/D734A^ data were collected with 0.1 mM RBD1^D623A/D734A^, 25 °C, and pH 7.0.

**Fig. S8.**
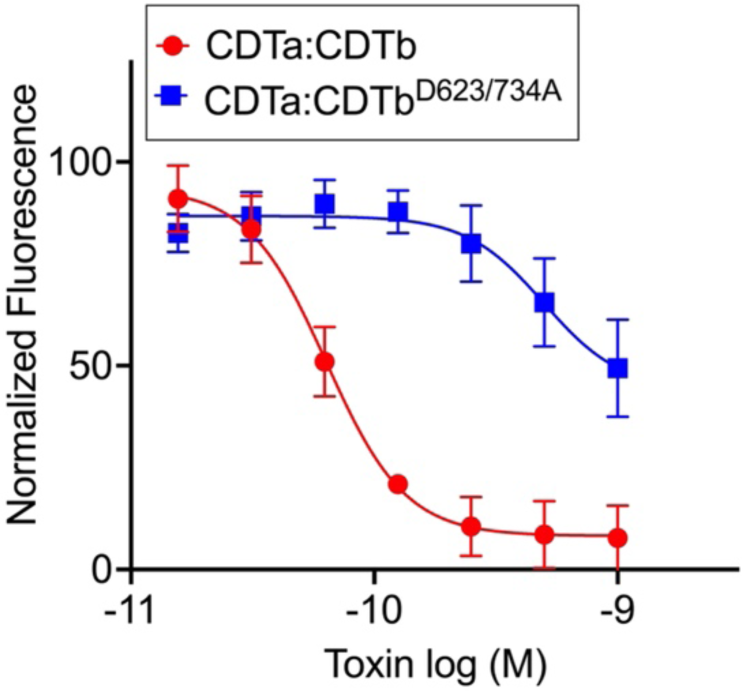
*Vero* cell toxicity from CDTa addition to either CDTb^WT^ (red, TC_50_ = 70 ± 20 pM) or CDTb^D623A/D734A^ (blue, TC_50_ = 560±60 pM). For simplicity, the *X*-axis is presented using the CDTa concentration, but each experiment contains a 7x concentration of activated CDTb subunits as described previously (6, 11).

**Fig. S9.**
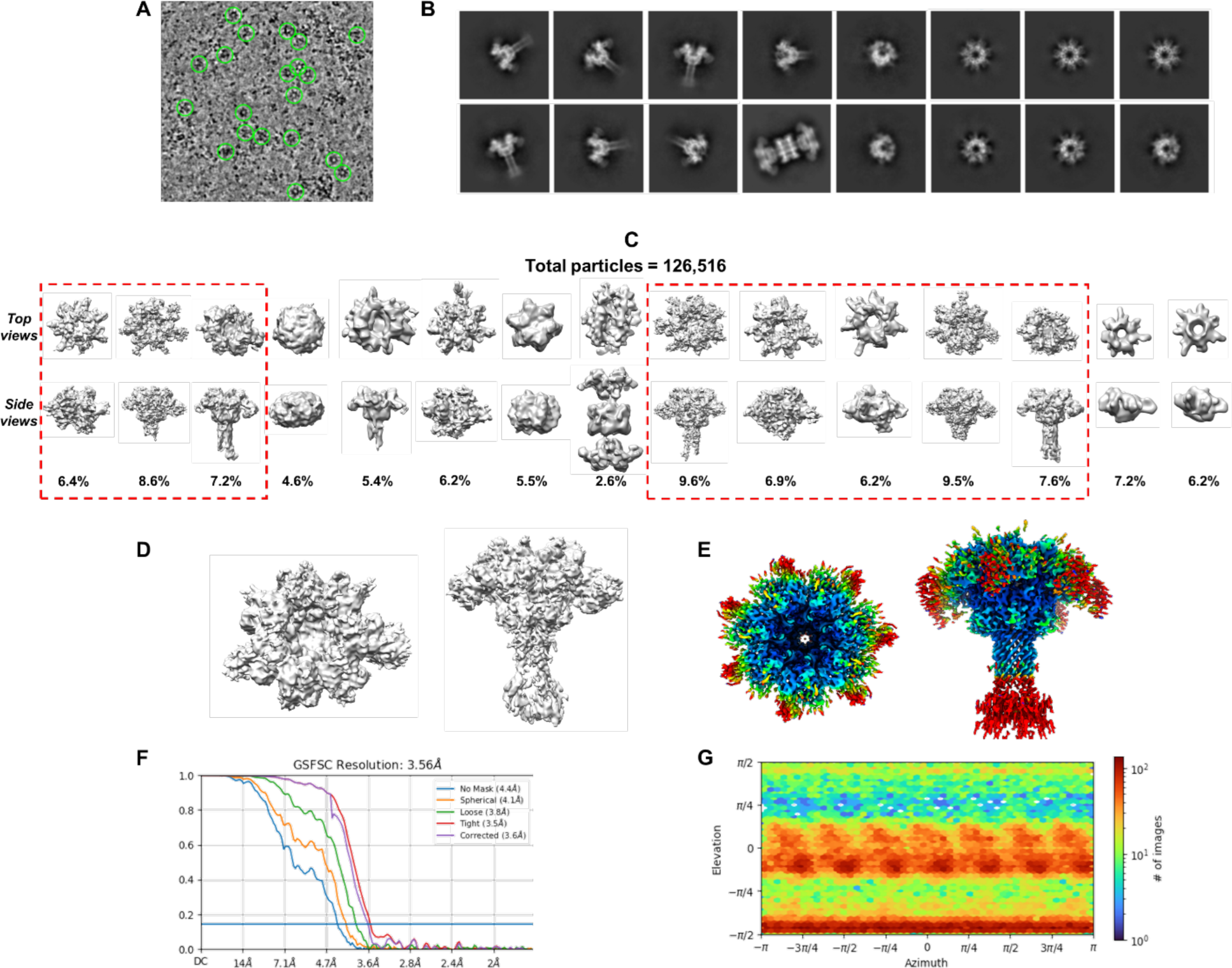
Structural characterization of CDTb^D623A/D734A^. (*A*) Example of a motion corrected micrograph with a few particles highlighted in circles to show different orientations. (*B*) Results of 2D classification generated by cryoSPARC. (*C*) Results of 3D classification generated by cryoSPARC. Classes that were combined in final reconstruction are highlighted. (*D*) Resulting electron density map at C1 symmetry. (*E*) Resulting electron density map colored by local resolution after application of C7 symmetry element. (*F*) FSC curve from the final reconstruction with C7 symmetry imposed. (*G*) Directional distribution of particle orientation.

**Fig. S10.**
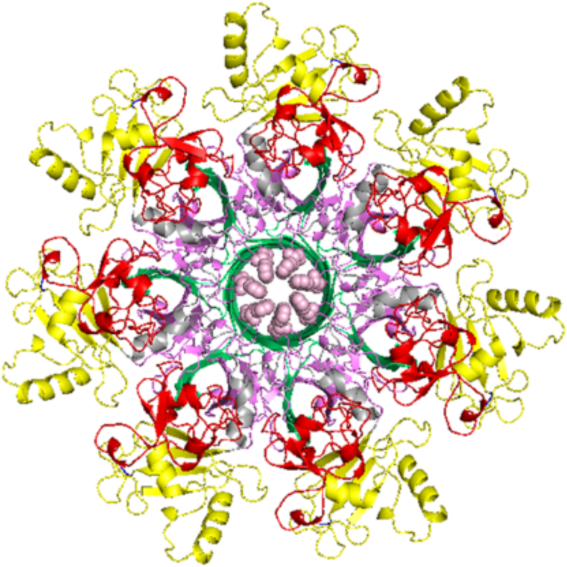
Top-view projection of the “φ-gate” assembly in the CDTb^D623A/D734A^ resembles the closed pore conformation identified in WT CDTb pore state structure. Phe455 residues comprising the pore in the mutant are shown in light pink. HD1, β-BD, HD2, and HD3 are shown in red, green, violet, and yellow, respectively.

**Fig. S11.**
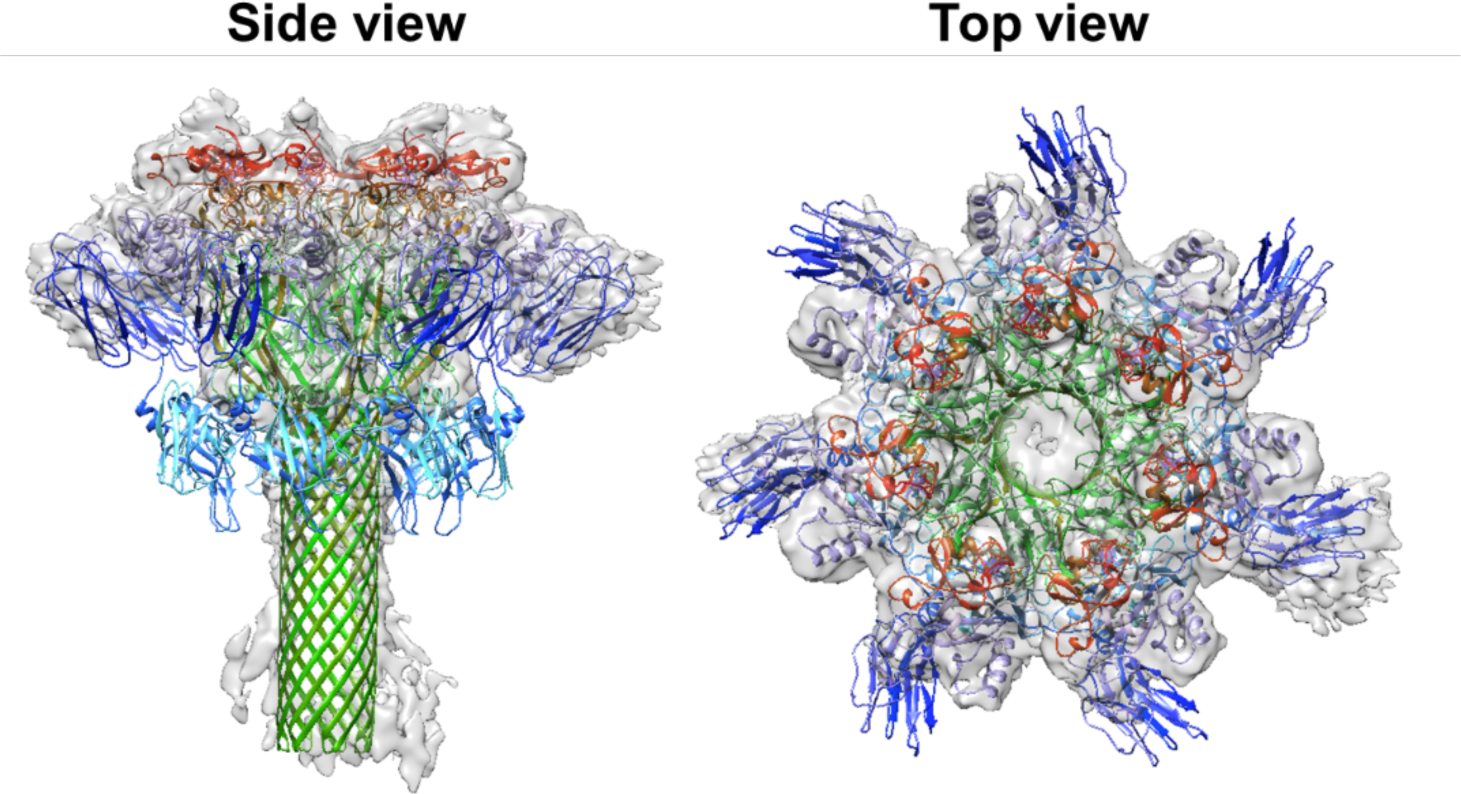
Superimposition of the heptamer of WT CDTb and electron density map of CDTb^D623A/D734A^ solved at C1 symmetry. Side and top view projections highlight the electron density of RBD1 (blue) in a single protomer. HD1, β-BD, HD2, HD3, RBD1, and RBD2 are shown in red, green, violet, yellow, blue, and cyan, respectively.

**Fig. S12.**
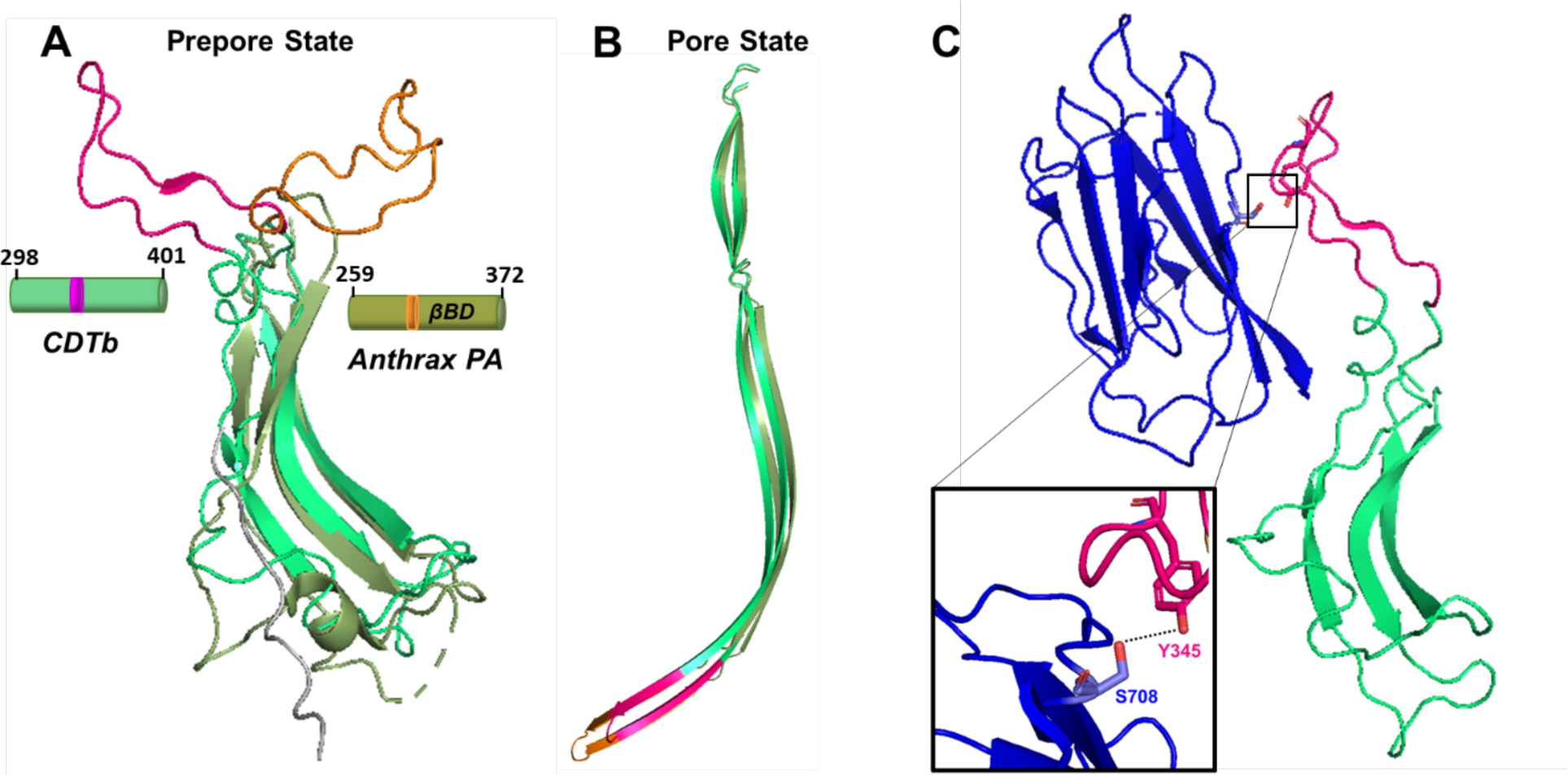
Comparison of β-barrel forming domains (βBD) orientations in protomers of CDTb and anthrax PA. (*A*) Superimposition of prepore state βBD of CDTb (lime green; residues 298-401 in PDB: 6UWT) and anthrax PA (olive green; residues 259-372 in PDB: 1TZO). β-loops forming the tip of the extended β-barrel in the pore state of each toxin are rotated towards opposite directions. In CDTb, the β-loop (magenta; residues 339-356) is projected towards the RBD1 and in anthrax PA, the corresponding loop (orange; residues 302-323) packs between the domain 2 and 4 of the neighboring protomer. *(B)* Superimposition of the pore states for βBD of CDTb (PDB: 6UWR) and for anthrax PA (PDB: 3J9C) depicts identical structural folds. The only notable difference is that βBD in anthrax PA is longer than that of βBD in CDTb. Residues forming the tip of the β-barrel in the pore state are oriented as a β-loop in the prepore state projecting towards opposite directions in both toxins. *(C)* The interface of βBD and RBD1 (blue; residues 616-744) in CDTb prepore state. Close-up view of the hydrogen bond interaction (dotted line) between Y345 of β-loop in βBD and S708 of RBD1 is highlighted in the inset figure.

**Table S1.**
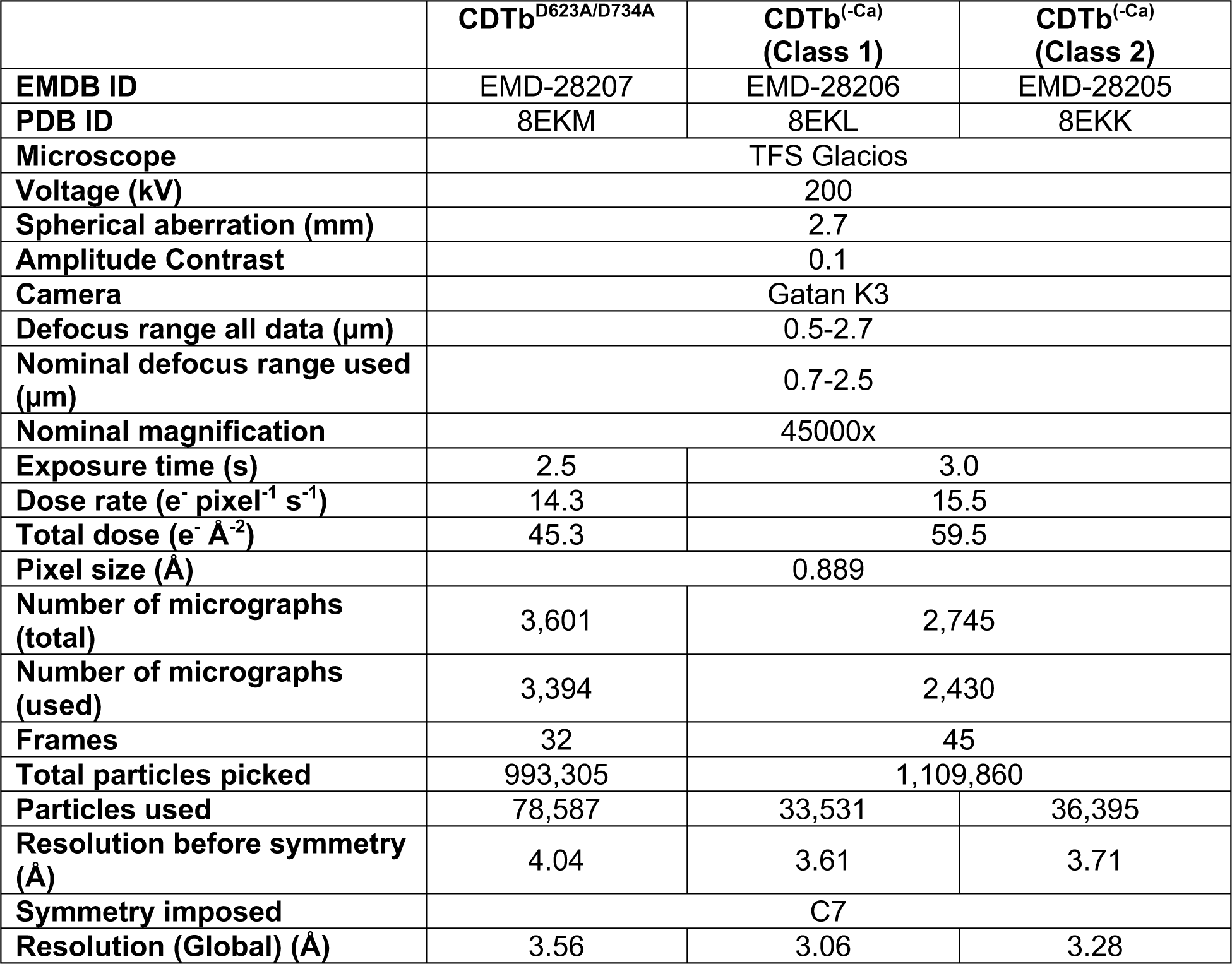
Details and parameters on cryoEM data collection.

## Notes

### Competing Interest Statement

The authors have declared no competing interest.

### Summary of Updates

Changes to manuscript title Updated order of authors Supplemental files updated

