## Supplemental Information for "Membrane binding and pore formation is Ca^2+^-dependent for the *Clostridioides difficile* binary toxin"

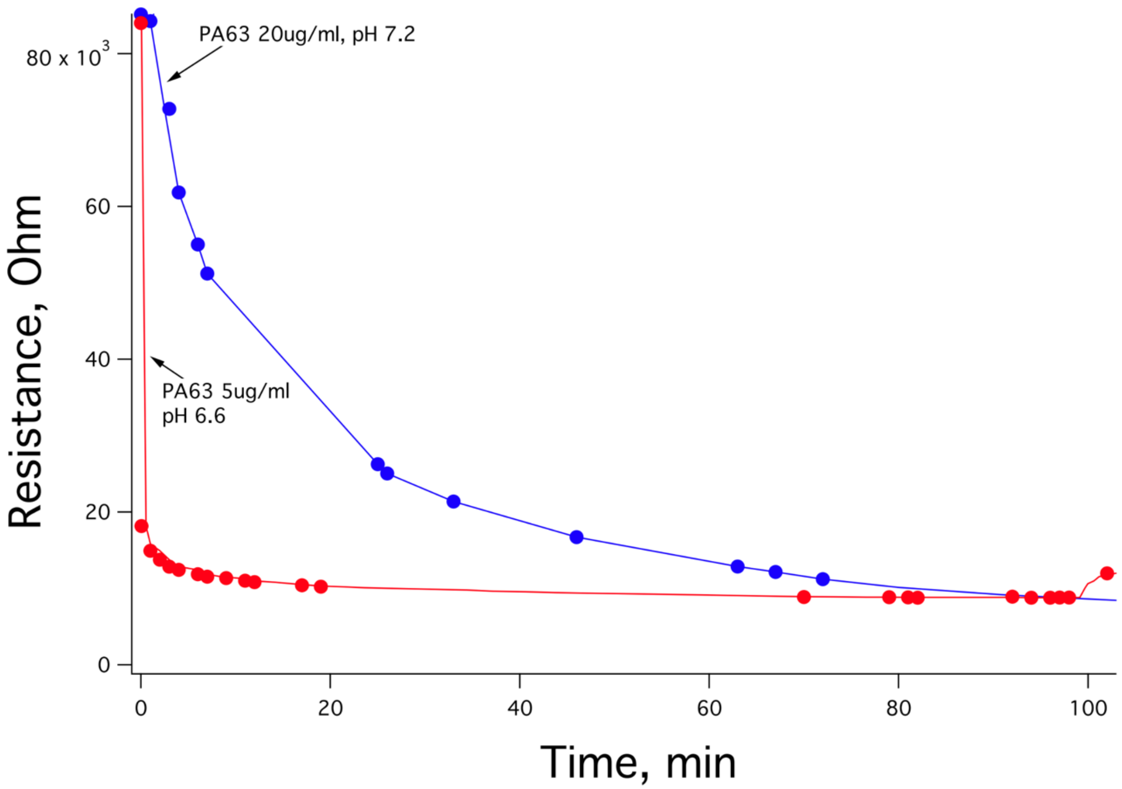


**Fig. S1.**
Membrane interactions of anthrax PA63 at pH 7.2 and pH 6.6. Resistance measured by electrochemical impedance spectroscopy shows rapid protein insertion into lipid membrane at lower pH *(23)*.

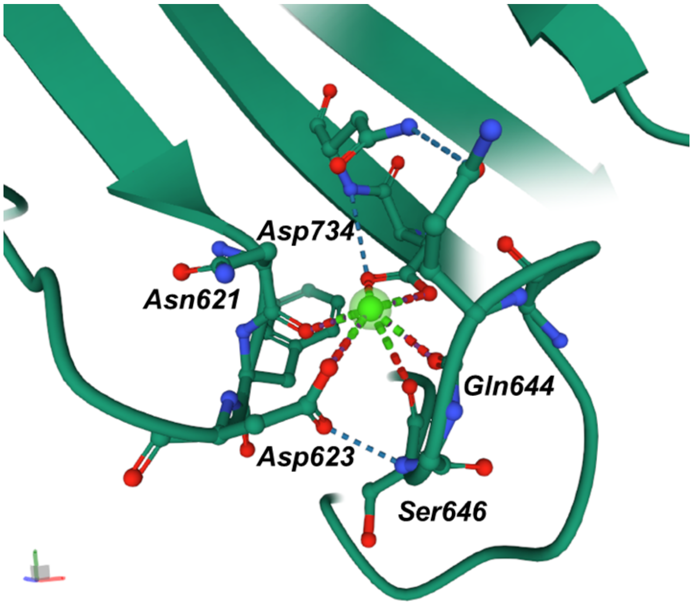


**Fig. S2.**

$Ca^{2+}$ binding site in RBD1. CDTb wild type (PDB: 6UWR) contains a single $Ca^{2+}$ ion (shown in bright green) in the RBD1 domain and the liganding oxygen atoms are from Asn621, Asp623, Ser646, Gln644, and Asp734. Point mutations were targeted at the two aspartic acid residues, Asp623 and Asp734, which were converted into alanine residues as discussed.

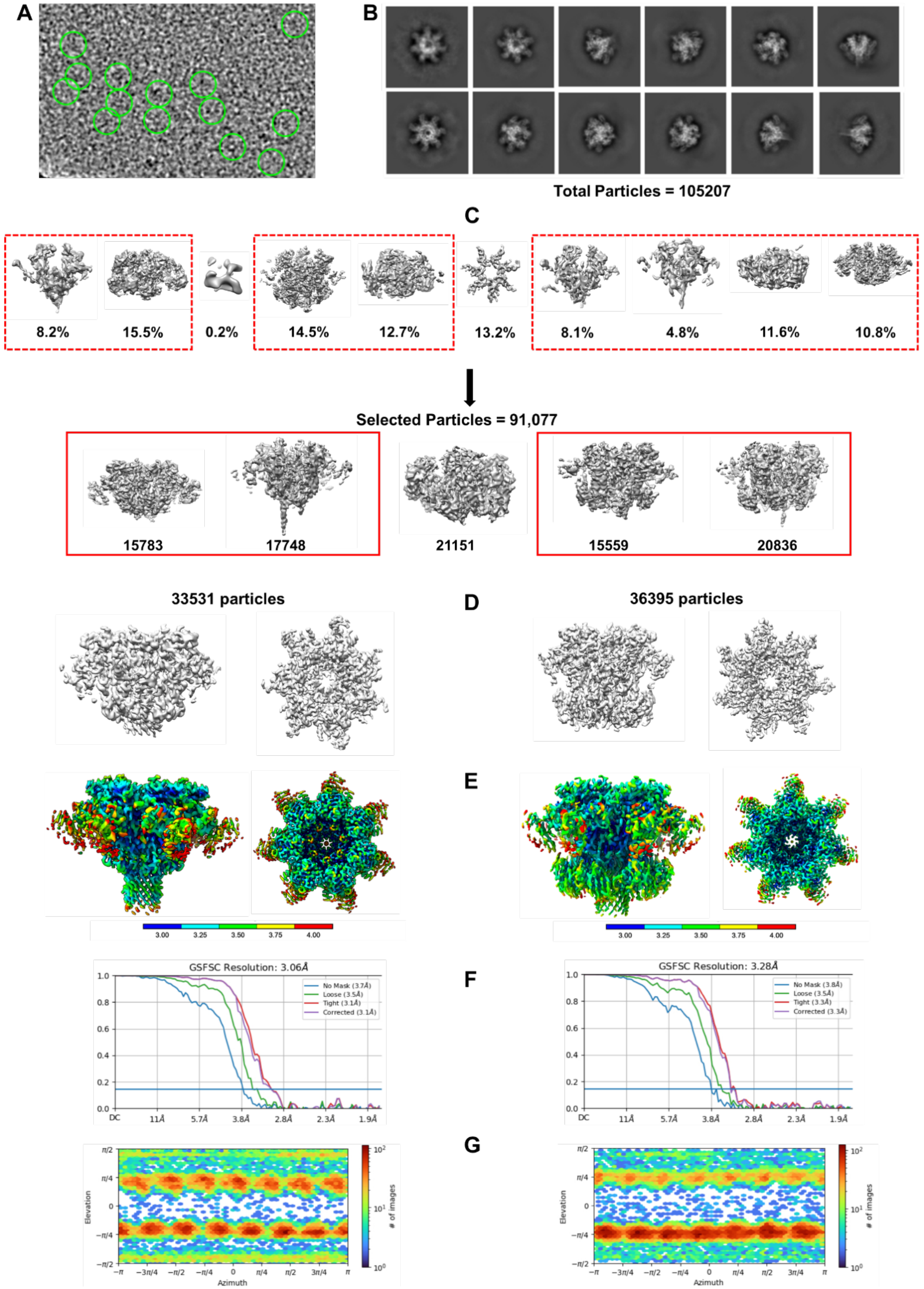
A
B
Total Particles = 105207
C
8.2%
15.5%
0.2%
14.5%
12.7%
13.2%
8.1%
4.8%
11.6%
10.8%
Selected Particles = 91,077
15783
17748
21151
15559
20836
33531 particles
D
36395 particles
E
F
GSFSC Resolution: 3.06Å
GSFSC Resolution: 3.28Å
G
Elevation
Azimuth
### of images

**Fig. S3.**

Structural characterization of $Ca^{2+}$ depleted WT CDTb. (*A*) Example of a motion corrected micrograph with a few particles highlighted in circles to show different orientations. (*B*) Results of 2D classification generated by cryoSPARC. (*C*) Results of 3D classification generated by cryoSPARC. Classes that were combined in the final reconstructions are highlighted in red boxes. RBD2 distorted structure (class 1) is shown on the left side. The RBD2 intact structure (class 2) is shown on the right side. (*D*) Resulting electron density maps at C1 symmetry. (*E*) Resulting electron density maps colored by local resolution after application of C7 symmetry element. (*F*) FSC curves from the final reconstruction with C7 symmetry imposed. (*G*) Directional distributions of particle orientation.

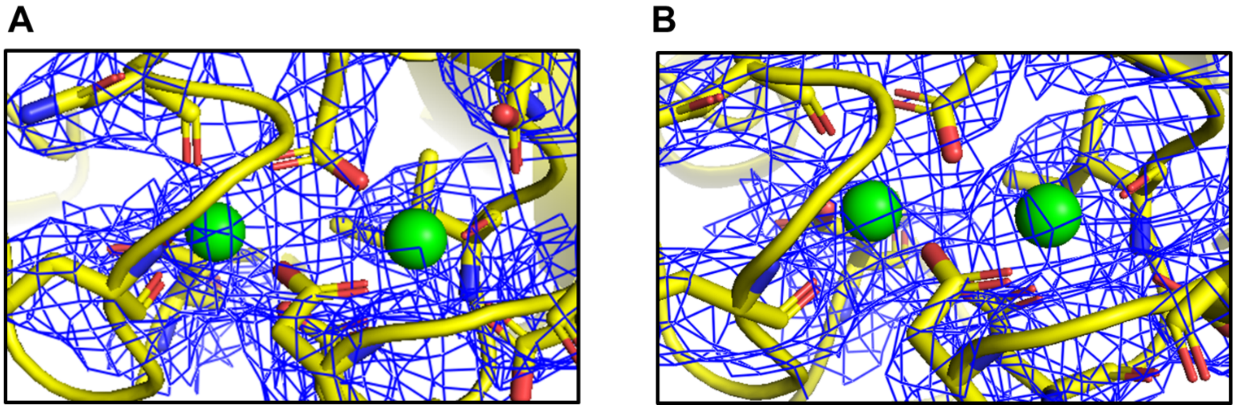


**Fig. S4.**

High affinity dual $Ca^{2+}$ binding sites located in the N-terminal heptamerization domain 1 (HD1) of $Ca^{2+}$-depleted WT CDTb. Dialysis of WT CDTb against the chelex-treated 15 mM HEPES buffer (pH 7.0), 150 mM NaCl, 2 mM EDTA, and 2 mM EGTA only removed the $Ca^{2+}$ in RBD1 while tightly bound $Ca^{2+}$ in HD1 remained intact. The coulomb potential maps (i.e., cryoEM electron isomesh maps; blue) in both classes of $Ca^{2+}$-depleted WT CDTb (*A*) class 1 with increased flexibility in both RBD1 and RBD2; (*B*) class 2 with increased flexibility in RBD1 only, depicted electron density for two $Ca^{2+}$ ions (Ca1, Ca2; green spheres) in the HD1. Calcium liganding residues identified involved oxygen atoms of D222/D224/E231/D273/N260/E263 for the Ca1 site, and D220/D222/D224/E321/D228/I226(C = O) for the Ca2 site.

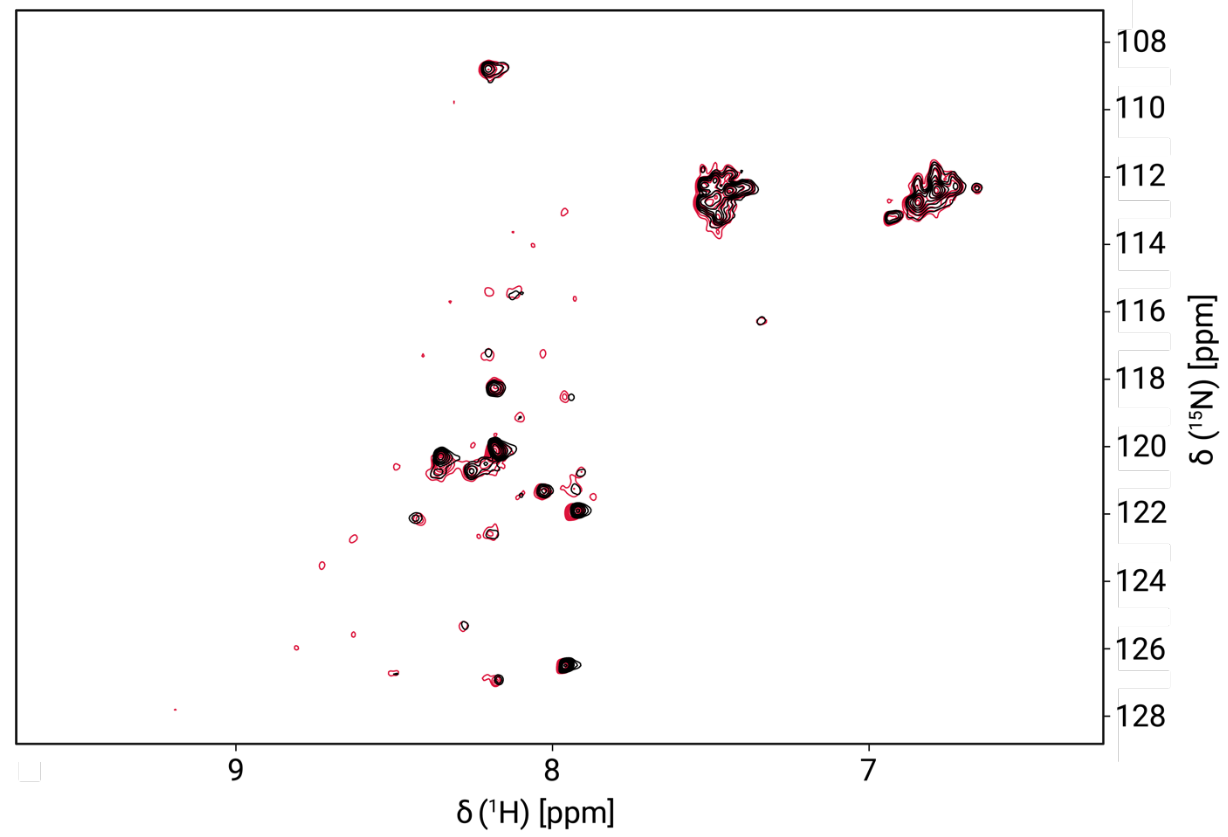


**Fig. S5.**

Overlay of 2D $^{15}$N-edited HSQC NMR spectra of RBD1$^{D623A/D734A}$ without $Ca^{2+}$ (black) and 20 hours after incubation with 10 mM $Ca^{2+}$ (red). RBD1$^{D623A/D734A}$ NMR data were collected with 0.1 mM RBD1$^{D623A/D734A}$, 25 °C and pH 7.0.

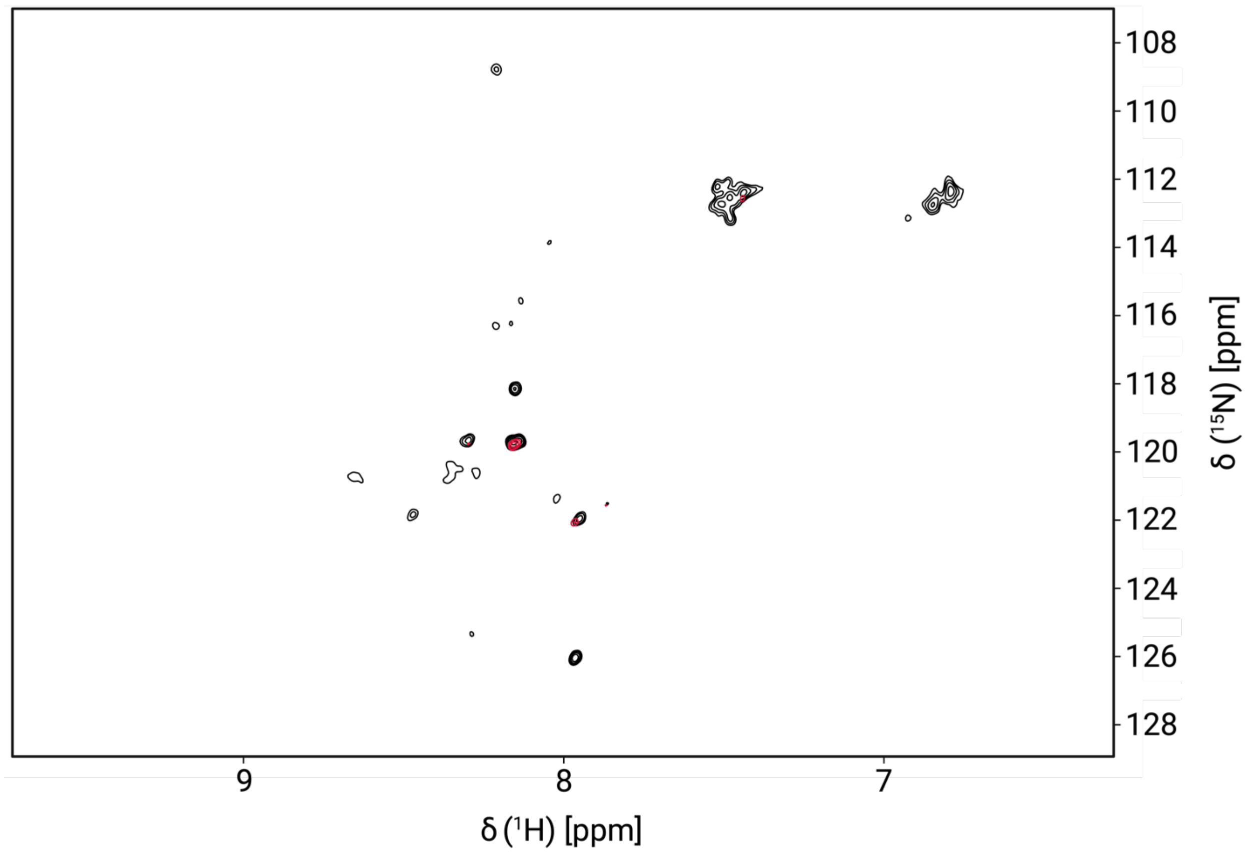


**Fig. S6.**
Overlay of 2D $^{15}$N-edited HSQC NMR spectra of RBD1$^{D623A/D734A}$ without $Ca^{2+}$ (black) and 20 hours after incubation with 10 mM $Ca^{2+}$ (red). RBD1$^{D623A/D734A}$ NMR data were collected with 0.1 mM RBD1$^{D623A/D734A}$, 25 °C, and pH 5.0.

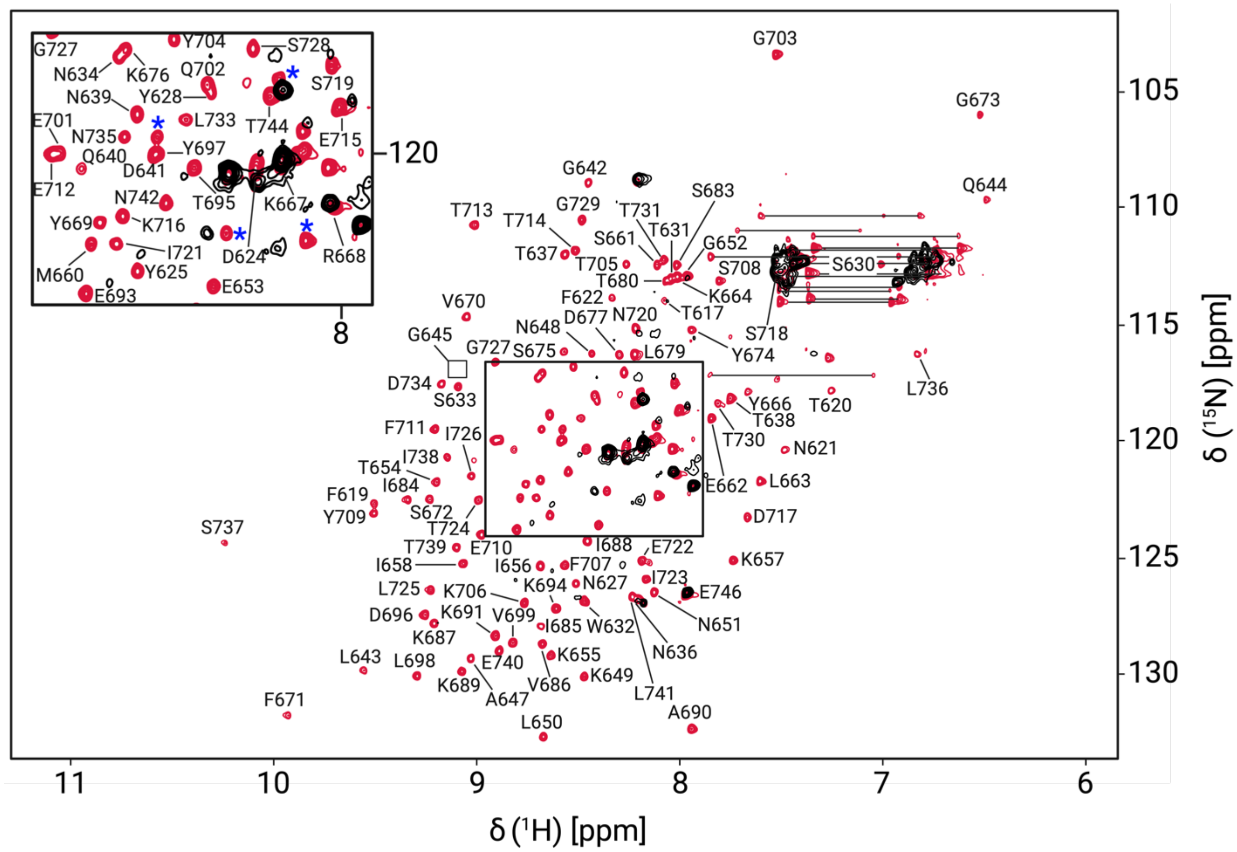


**Fig. S7.**

Overlay of 2D $^{15}$N-edted HSQC NMR spectra of RBD1$^{D623A/D734A}$ (black) and RBD1$^{WT}$ (red) after 20 hours incubation with 10 mM $Ca^{2+}$. Residues marked by blue asterisks are associated with His-tag. A correlation for G645 appeared in the noise and is indicated by a box. Correlations connected by horizontal lines correspond to sidechain $NH_2$ groups. RBD1$^{WT}$ data were collected with 0.25 mM RBD1$^{WT}$, 25 °C and pH 7.0; RBD1$^{D623A/D734A}$ data were collected with 0.1 mM RBD1$^{D623A/D734A}$, 25 °C, and pH 7.0.

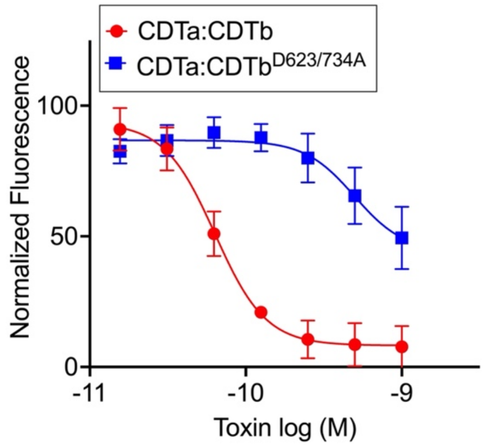


**Fig. S8.**

*Vero* cell toxicity from CDTa addition to either $CDTb^{WT}$ (red, $TC_{50} = 70 \pm 20$ pM) or $CDTb^{D623A/D734A}$ (blue, $TC_{50} = 560 \pm 60$ pM). For simplicity, the *X*-axis is presented using the CDTa concentration, but each experiment contains a 7x concentration of activated CDTb subunits as described previously (6, 11).

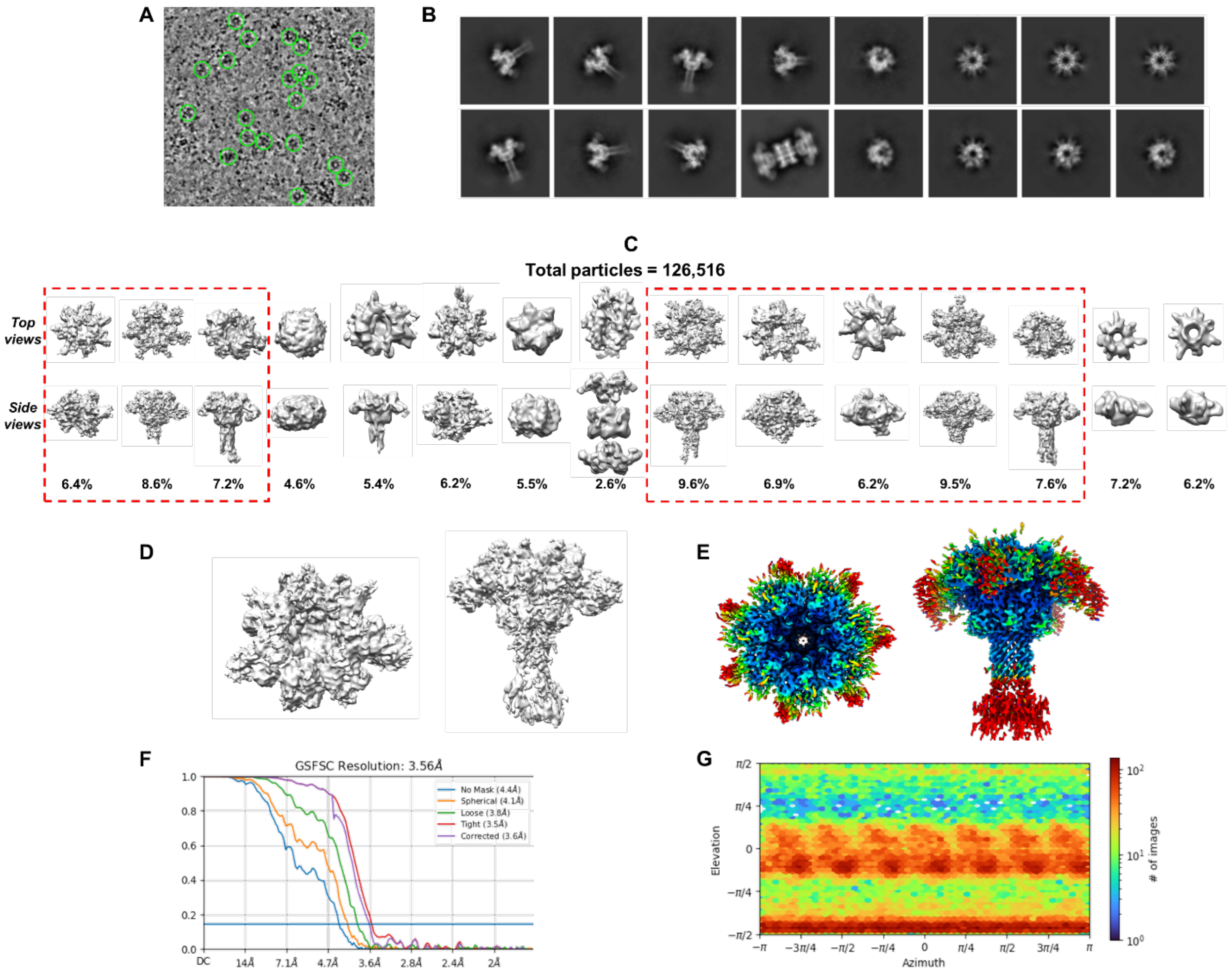


**Fig. S9.**

Structural characterization of CDTb$^{D623A/D734A}$. (*A*) Example of a motion corrected micrograph with a few particles highlighted in circles to show different orientations. (*B*) Results of 2D classification generated by cryoSPARC. (*C*) Results of 3D classification generated by cryoSPARC. Classes that were combined in final reconstruction are highlighted. (*D*) Resulting electron density map at C1 symmetry. (*E*) Resulting electron density map colored by local resolution after application of C7 symmetry element. (*F*) FSC curve from the final reconstruction with C7 symmetry imposed. (*G*) Directional distribution of particle orientation.

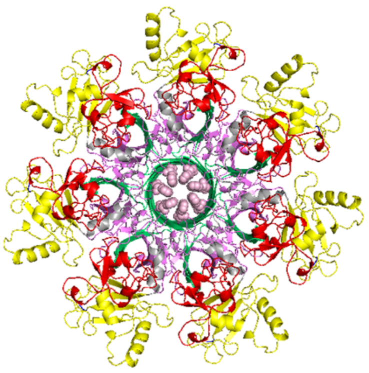

**Fig. S10.**

Top-view projection of the "φ-gate" assembly in the CDTb$^{D623A/D734A}$ resembles the closed pore conformation identified in WT CDTb pore state structure. Phe455 residues comprising the pore in the mutant are shown in light pink. HD1, β-BD, HD2, and HD3 are shown in red, green, violet, and yellow, respectively.

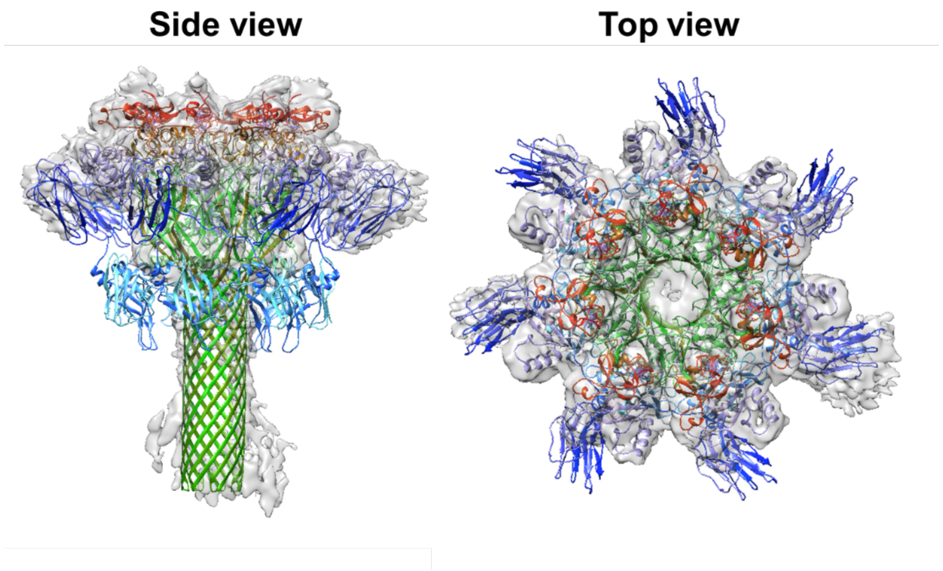


**Fig. S11.**
Superimposition of the heptamer of WT CDTb and electron density map of CDTb$^{D623A/D734A}$ solved at C1 symmetry. Side and top view projections highlight the electron density of RBD1 (blue) in a single protomer. HD1, β-BD, HD2, HD3, RBD1, and RBD2 are shown in red, green, violet, yellow, blue, and cyan, respectively.

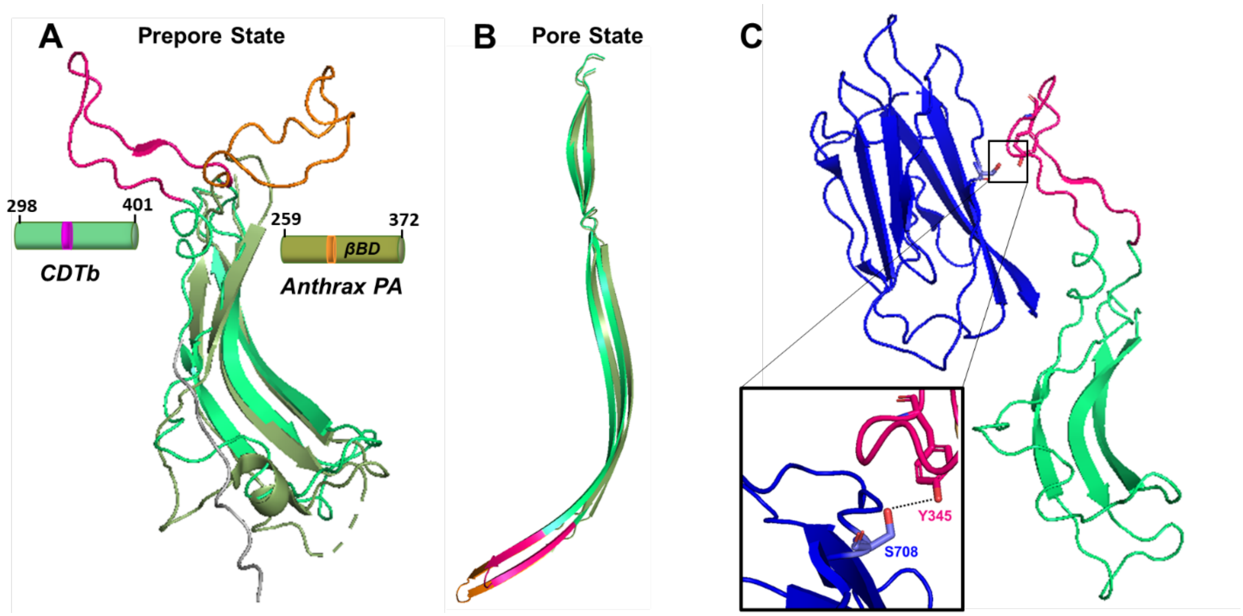


**Fig. S12.**
Comparison of β-barrel forming domains (βBD) orientations in protomers of CDTb and anthrax PA. (*A*) Superimposition of prepore state βBD of CDTb (lime green; residues 298-401 in PDB: 6UWT) and anthrax PA (olive green; residues 259-372 in PDB: 1TZO). β-loops forming the tip of the extended β-barrel in the pore state of each toxin are rotated towards opposite directions. In CDTb, the β-loop (magenta; residues 339-356) is projected towards the RBD1 and in anthrax PA, the corresponding loop (orange; residues 302-323) packs between the domain 2 and 4 of the neighboring protomer. *(B)* Superimposition of the pore states for βBD of CDTb (PDB: 6UWR) and for anthrax PA (PDB: 3J9C) depicts identical structural folds. The only notable difference is that βBD in anthrax PA is longer than that of βBD in CDTb. Residues forming the tip of the β-barrel in the pore state are oriented as a β-loop in the prepore state projecting towards opposite directions in both toxins. *(C)* The interface of βBD and RBD1 (blue; residues 616-744) in CDTb prepore state. Close-up view of the hydrogen bond interaction (dotted line) between Y345 of β-loop in βBD and S708 of RBD1 is highlighted in the inset figure.

| | CDTb$^{D623A/D734A}$ | CDTb$^{(-Ca)}$ (Class 1) | CDTb$^{(-Ca)}$ (Class 2) |
|---|---|---|---|
| **EMDB ID** | EMD-28207 | EMD-28206 | EMD-28205 |
| **PDB ID** | 8EKM | 8EKL | 8EKK |
| **Microscope** | TFS Glacios | | |
| **Voltage (kV)** | 200 | | |
| **Spherical aberration (mm)** | 2.7 | | |
| **Amplitude Contrast** | 0.1 | | |
| **Camera** | Gatan K3 | | |
| **Defocus range all data (µm)** | 0.5-2.7 | | |
| **Nominal defocus range used (µm)** | 0.7-2.5 | | |
| **Nominal magnification** | 45000x | | |
| **Exposure time (s)** | 2.5 | 3.0 | |
| **Dose rate (e$^-$ pixel$^{-1}$ s$^{-1}$)** | 14.3 | 15.5 | |
| **Total dose (e$^-$ Å$^{-2}$)** | 45.3 | 59.5 | |
| **Pixel size (Å)** | 0.889 | | |
| **Number of micrographs (total)** | 3,601 | 2,745 | |
| **Number of micrographs (used)** | 3,394 | 2,430 | |
| **Frames** | 32 | 45 | |
| **Total particles picked** | 993,305 | 1,109,860 | |
| **Particles used** | 78,587 | 33,531 | 36,395 |
| **Resolution before symmetry (Å)** | 4.04 | 3.61 | 3.71 |
| **Symmetry imposed** | C7 | | |
| **Resolution (Global) (Å)** | 3.56 | 3.06 | 3.28 |

**Table S1.**
Details and parameters on cryoEM data collection.
